# Rapid volumetric brain changes after acute psychosocial stress

**DOI:** 10.1101/2021.12.01.470604

**Authors:** Marie Uhlig, Janis D. Reinelt, Mark E. Lauckner, Deniz Kumral, H. Lina Schaare, Toralf Mildner, Anahit Babayan, Harald E. Möller, Veronika Engert, Arno Villringer, Michael Gaebler

## Abstract

Stress is an important trigger for brain plasticity: Acute stress can rapidly affect brain activity and functional connectivity, and chronic or pathological stress has been associated with structural brain changes. Measures of structural magnetic resonance imaging (MRI) can be modified by short-term motor learning or visual stimulation, suggesting that they also capture rapid brain changes. Here, we investigated volumetric brain changes (together with changes in T1 relaxation rate and cerebral blood flow) after acute stress in humans as well as their relation to psychophysiological stress measures.

Sixty-seven healthy men (25.8±2.7 years) completed a standardized psychosocial laboratory stressor (Trier Social Stress Test) or a control version while blood, saliva, heart rate, and psychometrics were sampled. Structural MRI (T1 mapping / MP2RAGE sequence) at 3T was acquired 45 min before and 90 min after intervention onset. Grey matter volume (GMV) changes were analysed using voxel-based morphometry. Associations with endocrine, autonomic, and subjective stress measures were tested with linear models.

We found significant group-by-time interactions in several brain clusters including anterior/mid-cingulate cortices and bilateral insula: GMV was increased in the stress group relative to the control group, in which several clusters showed a GMV decrease. We found a significant group-by-time interaction for cerebral blood flow, and a main effect of time for T1 values (longitudinal relaxation time). In addition, GMV changes were significantly associated with state anxiety and heart rate variability changes.

Such rapid GMV changes assessed with VBM may be induced by local tissue adaptations to changes in energy demand following neural activity. Our findings suggest that endogenous brain changes are counteracted by acute psychosocial stress, which emphasizes the importance of considering homeodynamic processes and generally highlights the influence of stress on the brain.

**Highlights:**

- We investigated rapid brain changes using MRI in a stress and a control group
- VBM-derived GMV showed a significant group-by-time interaction in several clusters
- Main pattern: GMV in the stress group increased relative to the control group, in which GMV decreased
- GMV changes across groups were associated with state anxiety and heart rate variability
- Neither cerebral blood flow, nor T1 values fully account for the VBM results

## Introduction

A stressor is a real or imagined threat to an organism’s integrity or well-being, which elicits a psychological and physiological stress response (Herman et al., 2003). Rapidly activated and rapidly terminated, the stress response is highly adaptive in situations of acute threat, but a chronically activated stress system can have detrimental effects and constitutes a major risk factor for physical and mental disease (McEwen & Gianaros, 2010). While the stress response is orchestrated by the brain, it involves the whole organism, particularly the autonomic nervous systm and endocrine systems, with the hypothalamic-pituitary-adrenal axis (HPA axis) as a central component (Kemeny, 2003). In turn, brain structure and function can be affected by stress, and brain plasticity associated with chronic stress has been detected with structural magnetic resonance imaging (MRI) (Spalletta et al., 2014). In the current study, we used structural MRI to investigate rapid brain changes after acute stress in humans.

The stress response comprises a cascade of hormonal signals including corticotropin-releasing hormone (CRH), vasopressin, adrenocorticotropic hormone (ACTH), and cortisol (Tsigos & Chrousos, 2002), which activates bodily functions to counteract the stressor. Most importantly, it triggers suppression of the immune system, faster glucose metabolization, and increased blood pressure (Cohen et al., 1991; Nesse et al., 2016). Being lipophilic, cortisol can cross the blood-brain barrier and, through its action on brain structures such as the hippocampus, terminate the stress response (Tasker & Herman, 2011; Joëls et al., 2013). This highlights the strong association of cortisol with long-term effects of stress on brain plasticity (McEwen & Gianaros, 2011), which occurs predominantly in regions involved in HPA axis regulation, such as prefrontal cortex (PFC), hippocampus, and amygdala (McEwen & Gianaros, 2011).

Brain plasticity describes the brain’s capacity to alter its structure and function to adapt to changing demands (Lövdén et al., 2010). Brain structure and function are thereby inseparable, with structure constraining function and function shaping structure In a supply-demand model, regional volume changes represent a continuous adaptation of the brain in supply (e.g., brain tissue) to changing environmental demands, mediated by alterations in activity (Lövdén et al., 2013). In support of this model, MRI studies often report a parallel development of structural and functional networks (He et al., 2007). Simulations suggest that the structure-function relationship is determined by biomechanical features, which are affected by hemodynamic processes following neural activity (Zoraghi et al., 2021). The strength of structure-function relationships thereby varies regionally: in sensory and unimodal regions, function may be more strongly constrained by structure than in transmodal regions like the cingulate cortex or the insulae (Valk et al., 2022), which are reportedly involved in stress processing, which also exhibit more synaptic plasticity (Mesulam et al., 1998).

Stress-induced functional brain changes have been shown using MRI: During stress, the BOLD signal increased in prefrontal areas (Dedovic et al., 2009; Wheelock et al., 2016) and decreased in subcortical regions, including the hippocampus (Dedovic et al., 2009; Pruessner et al., 2008). Such stress-related brain changes in the PFC and subcortical regions also outlasted the stress task, which was ascribed to sustained vigilance or emotional arousal (Wang et al., 2005). Stress-related changes in functional connectivity have been shown in the salience network (Hermans et al., 2014), including the anterior cingulate cortex (ACC) and other cortical midline structures (Veer et al., 2011). These stress-related functional connectivity changes also correlated with individual cortisol trajectories (Veer et al., 2012). In the framework by Hermans et al. (2014), humans adapt to acute stress by reallocating resources to brain networks that implement adaptive mental states: the salience network (associated with emotional reactivity, fear or vigilance) *during* and the executive control network (associated with higher-order cognition) *after* an acute stressor. When the stress wanes, the resource allocation to these two networks “normalizes” and with it the relative importance of emotional reactivity and higher-order cognition (Hermans et al., 2014). Analysing the resting-state fMRI from the experiment presented here, we previously found a stress-related increase in thalamic functional connectivity (part of the salience network), which was linked to subjective stressfulness (Reinelt et al., 2019).

The link between chronic or pathological stress and structural brain changes in humans has been well-established (for a review see Radley et al., 2015): For example, stress-related psychopathologies have been associated with structural plasticity mainly in limbic and prefrontal areas (McEwen, 2005). Patients with post-traumatic stress disorder (PTSD) showed decreased grey matter volume (GMV) in the hippocampus (Chen et al., 2006; Karl et al., 2006), amygdala and ACC (Karl et al., 2006; Rogers et al., 2009). Also without a clinical diagnosis, higher levels of self-reported chronic stress have been associated with lower GMV in the hippocampus, amygdala, insula, and ACC (Ansell et al., 2012b; Lotze et al., 2020; Papagni et al., 2011).

In animal models, rapid stress-induced structural brain changes that have been detected within hours after acute stress exposure include attenuation of neurogenesis (marmosets: Gould et al., 1998, rats: Heine et al., 2004), changes in astrocyte density (in degus; Braun et al., 2009) or decreases in dendritic spine density (in mice; Chen et al., 2010). In the latter study, a mediating function of the HPA axis in stress-induced memory deficits and associated brain structural changes was suggested.

Here, we used the Trier Social Stress Test (TSST, Kirschbaum et al., 1993), a strong and naturalistic psychosocial stressor in humans, and MRI to investigate rapid structural brain plasticity after acute stress.

While subtle processes like dendritic remodelling are unlikely to be captured with MRI at a voxel size of 1.5 mm, experience-induced brain changes have been detected with structural MRI in humans. Such brain changes are typically investigated using voxel-based morphometry (VBM; Ashburner & Friston, 2000; Draganski et al. 2004), which uses computational tissue classification based on T1-weighted images to detect differences in brain tissue composition. Numerous VBM studies have found rapid and spatially specific experience-induced brain changes: for example, increased GMV in the motor cortex was found after one hour of balance training (Taubert et al., 2016) and after one hour of brain-computer-interface training in targeted brain regions (Nierhaus et al., 2021). Even after less active interventions, such as ten minutes of high-frequency visual stimulation (Naegel et al. 2017), 263 seconds of passive image viewing (Månsson et al. 2020) or twelve minutes of finger tapping (Olivo et al., 2022), GMV changes were found with VBM. Most evidence for rapid MRI changes comes from studies that involve the acquisition of novel skills or exposure to novel stimuli.

A stressful experience contains memory and learning aspects, as stress influences memory formation (Schwabe et al., 2022) and is an important trigger for learning (e.g., to foresee and adaptively react to future stressors; Peters et al., 2017). Therefore, similar mechanisms may underlie stress-related brain changes and brain changes induced by other types of sensorimotor experiences.

The physiology behind VBM-derived GMV changes remains unclear^1^. Theoretically, genesis of neurons, glia cells, and synapses as well as vascular changes could underlie structural MRI changes in GM (Zatorre et al., 2012). As described above, in rodents, these plastic processes have been found after stressful interventions (Chen et al., 2010; Braun et al., 2009). Animal studies that combined MRI and histological examination after training interventions have suggested neural dendrites and astrocytes as drivers of rapid, experience-induced brain changes in structural MRI (Keifer et al., 2015; Sagi et al., 2012). Both can occur after minutes to hours (Johansen-Berg et al., 2012).

Rapid GMV changes may also occur with alterations in the participant’s physiological state during MRI, for example by changes in hydration (Streitbürger et al., 2012) or osmolality (Höflich et al., 2017; Streitbürger et al., 2012). Furthermore, vascular changes can impact VBM results, because blood and GM have similar T1 relaxation values at 3T (Tardif et al., 2017; Wright et al., 2008), and changes in blood oxygenation and tissue oxygenation (Tardif et al., 2017) or cerebral blood flow (CBF; Franklin et al., 2013; Ge et al., 2017) may influence changes in VBM-derived GMV.

Yet, several studies show rapid structural MR changes independent of vasculature-related changes (e.g., Zaretskaya et al., 2022, Olivo et al., 2022, Nierhaus et al., 2021), suggesting additional or alternative mechanisms of VBM changes (Nierhaus et al., 2021). Moreover, physical activity-induced (without learning or stimulation) CBF increases are not necessarily accompanied by morphological changes (Olivo). Taken together, experience-induced brain changes may rely on non-vascular processes (possibly related to glial processes) and be driven by novelty or learning aspects of the experience.

To specify the stress-related structural plasticity found with VBM and clarify the contribution of vasculature, we complemented VBM with other MRI measures: CBF measured with pulsed arterial spin labelling (pASL) and T1 mapping. An increase in T1 values reflects an increase in water content (Fullerton et al., 1982) and, in the context of training-induced plasticity, increased T1 values has been discussed to reflect an increase in vascular tissue (Thomas et al., 2018). On the other hand, increased oxygenation following a breathing challenge has been shown to decrease T1 values (Tardif et al., 2017), which has been ascribed to the so-called tissue oxygenation-level dependent (TOLD) contrast (Haddock et al. 2013). To investigate differences between T1 maps, T1-weighted images and (preprocessed) VBM images, we also analysed intensity values from (unpreprocessed) T1-weighted (UNI) images within the VBM clusters.

Not only interventions but also endogenous changes at different time scales can affect measures of GMV: Ageing is a strong predictor for GMV decreases (Karch et al., 2019), but rhythmic GMV changes have also been reported over the course of the menstrual cycle and its hormonal fluctuations (Barth et al., 2016; Lisofsky et al., 2015) or with the circadian rhythm (Karch et al., 2019; Nakamura et al., 2015; Orban et al., 2020; Trefler et al., 2016): Total GMV decreased linearly from morning to afternoon in several studies (Karch et al., 2019; Nakamura et al., 2015; Trefler et al., 2016), particularly in medial prefrontal areas and the precuneus (Trefler et al., 2016). In addition, CSF increased over the course of the day (Trefler et al., 2016) whereas total white matter decreased in one study (Trefler et al., 2016), but was not associated with time of day in another (Karch et al., 2019). The circadian system and the stress system both maintain homeostasis by adapting to environmental conditions, and they strongly interact on the physiological level, with the HPA axis being a major component of both systems (Nader et al., 2010; Nicolaides et al., 2014). Given this relatively new evidence for circadian brain changes, the majority of experiments on experience-induced plasticity did not control for time of day.

To summarize, rapid brain changes have been detected with structural MRI in humans upon exogenous stimulation and with endogenous fluctuations, and in animals following stress exposure. We hypothesized that acute stress, as a relevant exogenous stimulus triggering an endogenous process (i.e., the stress response), can induce rapid volumetric brain changes in GM derived from MRI. To test this hypothesis, we had young, healthy men complete either a psychosocial stress test (Trier Social Stress Test, TSST; Kirschbaum et al., 1993) or a closely related control intervention without the psychosocially stressful component (placebo-TSST; Het et al., 2009). Before and after the intervention, we acquired MRI data. Throughout the entire experiment, we regularly sampled autonomic, endocrine, and subjective markers of the stress response (Figure 1). This enabled us to measure the stress response on three levels and assess their relationships with brain changes. We thereby expected brain changes to be more pronounced in subjects with stronger increases in autonomic, endocrine, and subjective stress measures.

**Figure 1.**
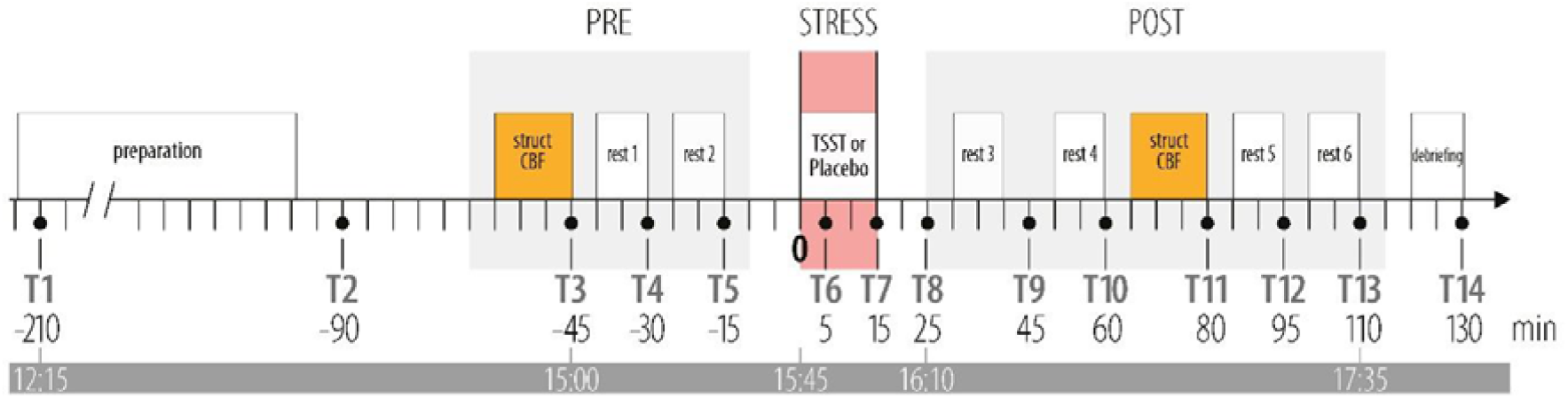
Schematic overview of the experiment: Between-subject design with the stress group (n = 32) undergoing the Trier Social Stress Test (TSST) and the control group (n = 34) a placebo-TSST. Orange boxes indicate two structural scans (struct) using T1 mapping (MP2RAGE) and two scans of pulsed arterial spin labelling for imaging cerebral blood flow (CBF). Psychometric ratings, saliva samples, and blood samples were acquired at 14 time points throughout the experiment (T1-T14). Minutes are relative to the onset of the intervention (TSST or placebo-TSST), while the bottom bar informs about the time-of-day. The grey boxes indicate phases in the MRI. (The TSST and the placebo-TSST took place outside of the MRI.)

As stress-induced brain changes have often been reported in the amygdala and the hippocampus (see above), they served as regions-of-interest (ROIs), complemented by an exploratory whole-brain analysis. To better depict the physiology of stress-induced brain changes, we also compared CBF and T1 values before and after the intervention. Additionally, we investigated the relation between GMV changes and the other (i.e., autonomic, endocrine, and subjective) stress measures.

## Methods

### Participants

We recruited male participants between 18 and 35 years of age via leaflets, online advertisements, and the participant database at the Max Planck Institute for Human Cognitive and Brain Sciences in Leipzig. Exclusion criteria, as assessed in a telephone screening, were smoking, excessive alcohol / drug consumption, past or current participation in psychological studies, regular medication intake, history of cardiovascular or neurological diseases, and a BMI higher than 27. In addition, standard MRI exclusion criteria applied, such as tattoos, irremovable metal objects (e.g., retainers, piercings), tinnitus, and claustrophobia.

We tested 67 young, healthy males. Because of an incidental medical finding, one participant was excluded, so that 66 participants (age: 25.8 ± 2.7, 21-32 years) entered the analyses, 32 in the stress and 34 in the control group.

On separate days prior to the stress/control paradigms as part of a separate study (Babayan et al., 2019), participants underwent extensive baseline measurements that included cognitive testing, blood screening, anthropometrics, structural and resting-state functional MRI scans, resting-state electroencephalography (EEG), self-report questionnaires, and a structured clinical interview (for details, see Babayan et al., 2019). If exclusion criteria were detected during the baseline assessment, participants were excluded from further testing.

Included participants were randomly assigned to either the stress or the control group. To avoid experimenter biases, the administrative staff remained blind to the testing condition until the first MRI session. All appointments were scheduled for the same time of day (11:45 am) to control for diurnal fluctuations of hormones (e.g., cortisol and ACTH; Nader et al., 2010; Nicolaides et al., 2014). Participants were asked to sleep at least 8 hours in the night before the experiment, to get up no later than 9 am, have a normal breakfast and then to not eat or exercise until their study appointment while also refraining from drinking coffee, black tea, or other stimulant drinks. Written informed consent was obtained from all participants. The study was approved by the ethics committee of the medical faculty at Leipzig University (number 385-1417112014), and participants were financially compensated.

Stress and the control groups did not differ significantly in age, hours of sleep on the day of testing, average sportive activity per week, or self-reported chronic stress (Reinelt & Uhlig et al., 2019).

### Procedure

The pre-scan was completed on average 45 (SD: ±3.9) min before intervention onset (before two resting-state fMRI scans, see Figure 1), and the post-scan was completed on average 88 (SD: ±3.6) min after intervention onset (between four resting-state fMRI scans, see Figure 1).

### Intervention

Each participant completed either a psychosocial stress test (Trier Social Stress Test, TSST; Kirschbaum et al., 1993) or the placebo-TSST as control intervention, which tightly controls for physical and cognitive load during the TSST (Het et al., 2009).

Participants in the stress group prepared for (5-min) and completed a job interview (5-min) as well as a difficult mental arithmetic task (5-min) in front of a committee (one female, one male professional actor), introduced as two professional psychologists trained in the analysis of nonverbal communication. Additionally, the task was recorded by a video camera and microphone. In the control condition, participants prepared (5-min) and spoke about their career aims (5-min) and solved an easy mental arithmetic task (5-min) with nobody else in the room and no video or audio recording. To extend the stressfulness of the TSST, participants in the stress group were told that a second task would follow during the scanning procedure. To make this scenario more plausible, participants were brought back to the scanning unit in the company of the experimenter and the TSST committee members. After rest 4, before the structural scan (+60 min after TSST onset), they were told that no additional task would follow. For a more detailed description of the interventions, see supplementary material (section 1.1. and 1.2.) and Reinelt & Uhlig et al. (2019).

Throughout the experiment, blood was sampled at 14 time points, and saliva and subjective ratings at 15 time points. At each sampling point, participants completed psychometric questionnaires, while autonomic and endocrine data were acquired. For further details, see below as well as Reinelt & Uhlig et al. (2019) and Bae & Reinelt et al. (2019).

### Magnetic resonance imaging

#### Acquisition

MRI was performed on a 3T MAGNETOM Verio (Siemens Healthineers, Erlangen, Germany) scanner with a 32-channel head-coil. The MP2RAGE sequence was used to acquire structural MR images. The MP2RAGE sequence yields a nearly bias-free T1-weighted (UNI) image, which is created by combining the two inversion images (INV1, INV2) and it produces a T1 map (T1) (Marques et al., 2010). The high-resolution MP2RAGE sequence had the following parameters (Streitbürger et al., 2014): TI1 = 700 ms, TI2 = 2500 ms, TR = 5000 ms, TE = 2.92 ms, FA1 = 4°, FA2 = 5°, 176 slices, voxel dimensions = 1 mm isotropic.

Cerebral blood flow (CBF) was measured using the pulsed arterial spin labelling (pASL) sequence implemented by the vendor (PICORE; Wong et al., 1997; Luh et al., 1999). For a detailed description of the pASL data acquisition, preprocessing, analysis and results, see supplement (section 1.4.).

#### Preprocessing

##### VBM

For each scan (pre-intervention, post-intervention), a brain mask was created from the INV2-images to remove the noisy background of the UNI images, which is a by-product of the division of the two inversion images. These background-masked T1-weighted images were preprocessed using the longitudinal preprocessing pipeline (with default settings, Version 1450 (CAT12.6) 2019-04-04) of the CAT12 toolbox (http://www.neuro.uni-jena.de/cat/) including intra-subject realignment, bias correction, segmentation into three tissue types (grey matter, white matter, and cerebrospinal fluid) and non-linear spatial registration to MNI space using DARTEL (Ashburner, 2007). By default, the images are resampled to a voxel size of 1.5 mm (isotropic) during preprocessing. We chose a resolution of 1.5 mm because it matches the resolution of the standard template used for registration and thereby avoid an additional interpolation step. Finally, the images were smoothed with a Gaussian kernel at 8-mm full-width at half maximum (FWHM). For further analysis, the segmented GM images were used.

##### T1 & T1-weighted

The background-masked T1-weighted images were warped to MNI space (using the *normalize:estimate&write* function in SPM). T1 maps were normalized to MNI space by applying the deformations from the normalization of the T1-weighted images. The normalized T1-weighted and T1 images were masked with the same sample-specific GM mask, which was used for the VBM analysis before smoothing with an 8-mm FWHM Gaussian kernel.

##### CBF

PASL time series were first realigned with FSL *McFlirt*, then normalized to MNI space using SPM12, and finally smoothed with a 3D spatial Gaussian filter (for details, see supplement, section 1.4.).

#### Postprocessing

For post-hoc analyses (see below for details), significant (p_FWE_ < 0.05) VBM clusters were saved as binarized NIfTI images from the result GUI and used as masks for post-hoc tests and to investigate changes in T1 and T1-weighted intensity values as well as CBF. For binarizing masks, multiplication with masks and extraction of GMV, T1 and T1-weighted intensity values, FSL was used (*fslmaths & fslstats* in *fslutils*, (Jenkinson et al., 2012).

##### VBM post-hoc

GMV values were extracted by multiplying binary masks of VBM clusters with the smoothed, preprocessed GM images and extracting the average value from each cluster.

##### T1 & T1-weighted

T1 and T1-weighted images were smoothed after applying a GM mask (at GM threshold 0.1). Values were extracted by multiplying binary masks of VBM clusters with the smoothed and normalized T1 and T1-weighted images and extracting the average value from each cluster. Additionally, average values from GM voxels outside of the VBM clusters were extracted to serve as a reference for potential global changes in T1 values or T1-weighted intensity values. Therefore, the smoothed, GM masked images were multiplied with an inverse binary VBM-cluster mask.

##### CBF

For the CBF analysis, the masks were resampled to a 2-mm isotropic voxel size to match the pASL images using the *coregister:reslice* function in SPM12. The preprocessed CBF maps were multiplied with binary masks for VBM clusters and the average CBF value for each cluster was extracted. As the pASL data is acquired within a manually defined slab, not all VBM clusters were fully covered (see Figure S2). Only clusters in which CBF values were available for at least 70% of voxels were included in the post-hoc CBF analysis.

#### Anatomical regions-of-interest definition

To test our regional hypotheses, anatomical regions-of-interest (ROIs) were created as binary masks of hippocampus and amygdala using the Anatomy toolbox (Eickhoff et al., 2005) and resampled to 1.5-mm space using SPM12 to match the anatomical images. ROI values were extracted by multiplying masks with the smoothed, modulated, warped, coregistered images using FSL (*fslmaths & fslstats* in *fslutils*, Jenkinson, et al., 2012). Below, “Hippocampus” and “Amygdala” (with capitalized first letters) refer to these anatomical ROIs.

#### Quality assessment

Image quality was assessed using the noise-to-contrast ratio (NCR), a quality parameter computed by the CAT12 toolbox from noise, bias and white-matter hyperintensities. Based on within-sample comparisons, data from participants whose image quality (NCR) was more than 3 standard deviations (SD) below the sample mean were excluded (see supplement, section 1.3. and Figure S3). Systematic changes in image quality were tested with a linear mixed model, which showed a significant group-by-time interaction effect for NCR (X^2^(1) = 7.9; p = 0.0049), driven by a significant decrease in image quality in the control group (t-ratio = 3.7, p = 0.0005). Head movement can negatively influence image quality in MRI (Power et al., 2015) as well as estimates of GMV and cortical thickness (Reuter et al., 2015). As no information about head movement was available from the MP2RAGE data, we calculated mean framewise displacement (MFD) as the sum of the absolute values of the six realignment parameters (Power et al., 2015) from the resting-state fMRI scans that directly preceded the MP2RAGE scan. Accounting for head motion by including MFD into the above model weakened the group-by-time interaction effect for NCR (X^2^(1) = 3.6; p = 0.059). Furthermore, a non-significant trend for an effect of MFD on NCR (X^2^(1) = 3.4, p = 0.068) was found, suggesting an association between the two quality parameters. To avoid circular analyses (since NCR was derived from the data), we included MFD in our statistical models to account for quality changes on volume estimates. MFD thereby served as a proxy covariate for image quality only. No participants were excluded based on motion parameters, but instead by using the CAT12 toolbox’s quality parameter “noise-to-contrast ratio” for detection. For extracted T1 and T1-weighted intensity values, values outside the range of 3 SD above and below sample mean were excluded (for details, see respective section below).

The quality assessment of CBF data is described in the supplement (section 1.4.3.).

### Psychophysiological stress measures

#### Autonomic

Heart rate (HR) and heart rate variability (HRV) were analysed from recordings of electrocardiography (ECG, outside MRI) and photoplethysmography (PPG, inside MRI). A detailed description of autonomic data acquisition and data preprocessing can be found in Reinelt & Uhlig et al. (2019). Autonomic recordings were binned into three-minute intervals. The average interbeat interval (the inverse HR) was determined for each interval and HRV was quantified as the square root of the mean squared differences of successive differences (RMSSD) in interbeat intervals, indexing parasympathetic cardio-regulation (e.g., Malik et al., 1996).

#### Endocrine

Blood and saliva samples were acquired throughout the entire experiment (inside as well as outside the scanner, see Figure 1). Saliva was sampled with a Sarstedt Salivette (Sarstedt, Nümbrecht, Germany) for at least 2 min per sample. Blood samples (serum and plasma; Sarstedt Monovette) were acquired by the experimenter from an intravenous catheter in the left or right cubital vein. Saliva and blood samples were analysed using Liquid chromatography-tandem mass spectrometry (LC-MS/MS) at the Institute for Laboratory Medicine, Clinical Chemistry and Molecular Diagnostics, University of Leipzig, following the protocol described in (Gaudl et al., 2016). A detailed analysis of changes in endocrine markers and their timing in the current study can be found in (Bae et al., 2019). For the present analysis, saliva cortisol and plasma ACTH were used to assess the association of GMV changes with endocrine stress measures at different times of HPA axis activation: ACTH, which is secreted earlier during HPA axis activation, peaked at 15 min after stressor onset, while saliva cortisol, a particularly robust stress marker (Vining et al., 1983), peaked at 25 min after stressor onset (see Bae et al., 2019; Reinelt & Uhlig et al., 2019). Participants with a cortisol increase below 1.5 nmol/l following psychosocial stress exposure can be considered non-responders and are often excluded from analyses including endocrine data (Miller et al., 2013).

#### Subjective

We presented questionnaires with OpenSesame 3.1.2 (Mathôt et al., 2012)□ on a laptop (outside MRI) or on a screen (inside MRI). Participants answered the questions with two keys on the laptop keyboard (outside MRI) or on an MRI-compatible button box (inside MRI). We here assessed state anxiety with the state trait anxiety questionnaire (STAI, sum score of the state subscale; (Laux, 1981; Laux & Spielberger, 2001) and the perceived stressfulness with the question “How stressed do you feel right now?”, which was answered using a visual analogue scale (VAS) with a sliding bar from 0 (“not at all”) to 100 (“very much”).

### Statistical Analysis

#### Analysis of neuroimaging data

For an illustration of the analysis pipeline see Figure S1 and S3.

#### Whole-brain analysis in SPM

Following quality assessment, three participants (two in the stress group) were excluded from the VBM analysis because of an NCR value more than 3 SD below the sample mean. The final VBM sample therefore consisted of 63 participants, 30 in the stress group and 33 in the control group. For statistical analysis of MRI data, delta images were created by subtracting the pre-intervention image from the post-intervention image. A two-sample t-test was performed on the difference images to model the group-by-time interaction. To focus the analysis on GM, thresholding is typically used in VBM analyses (e.g., Streitbürger et al., 2012). Since the voxel values in delta images describe a difference rather than the tissue probability itself, they could not be thresholded. Instead, we used a sample-specific GM mask. This mask is automatically created during model estimation in SPM; in our case a one-sample t-test on all smoothed, segmented GM images while applying an absolute masking threshold of 0.1 (probability of this voxel being GM) as recommended in the CAT12 manual (Version 30-06-2021, http://www.neuro.uni-jena.de/cat12/CAT12-Manual.pdf).

The total intracranial volume (TIV) was estimated for both images (pre-intervention, post-intervention) of each subject using CAT12, and their average was included as a covariate. To account for potential systematic, group-specific changes in image quality (see the section “Quality assessment” above and supplement, section 1.3.), MFD was included as a proxy for head motion as an additional covariate. The results from the two-sample t-test on Δgrey matter images (ΔGM) were investigated using two-sided t-contrasts (i.e., control > stress [1 -1 0 0] and control < stress [-1 1 0 0] with TIV and MFD in columns 3 and 4).

To minimize false positive and false negatives results, we used whole-brain threshold-free cluster enhancement (TFCE), a non-parametric multiple-comparison correction that does not require a cluster extent threshold, using the TFCE toolbox (http://dbm.neuro.uni-jena.de/tfce) with the default settings of 5000 permutations and the Smith-permutation method. Anatomical labels for significant clusters were found using the DARTEL-based “neuromorphometrics atlas” provided with the CAT12 toolbox. Below, we use capitalization to indicate the extracted anatomical labels (e.g., “Superior Medial Frontal Gyrus”)

#### Analysis of extracted imaging markers

Statistical analysis at the ROI level was performed using R 3.0.2 (R Core Team (2013); http://www.R-project.org/). Group differences in variables-of-interest over time were investigated with linear mixed models (LMMs; using the *lme4* package; (Bates et al., 2015), which included a random intercept for each subject to account for inter-individual differences. Visualizations were created in R using *ggplot* (Wickham, 2009) and by adapting raincloud plots (Allen et al., 2019).

#### Linear mixed model design

Across all analyses, the model was built following the same procedure (the full scripts can be found on https://gitlab.gwdg.de/necos/vbm.git):

1. A null model including a random intercept, covariates of no interest, as well as reduced fixed effects was set up and compared to a full model, which enables the targeted testing of effects of interest (Forstmeier & Schielzeth, 2011).
2. The full model was identical to the null model except for the effect of interest, in most cases the group-by-time interaction or other (i.e., autonomic, endocrine, or subjective) stress measures when testing their associations with GMV.
3. The difference between the full and the null model was tested using the *anova* function and setting the argument *test* to “chisq” to do a X^2^ (Chi^2^) test.
4. The *drop1* function was used to extract the results from the individual effects.
5. Non-significant interactions were dropped from the full model to reduce complexity (reduced model).
6. In case of significant interactions, the effects at the individual levels of predictors (e.g., within-group or for each cluster) were analysed post-hoc using the *emmeans & contrast* function with *Holm* correction from the *emmeans* package (Lenth, 2021). Estimated marginal means and 95% confidence intervals obtained with *emmeans* were used for plotting.

We tested the assumptions for LMMs by visually inspecting the distribution of residuals in a QQplot and a scatterplot of the residuals plotted against fitted values. The main criterion for the latter was symmetry along the y-axis. We also visually inspected residual plots for influential cases. Every case of excluded data is reported in the methods section and can be reproduced with the analysis scripts at https://gitlab.gwdg.de/necos/vbm.git. Multicollinearity was tested by extracting the variance inflation factor (VIF), using the *vif* function in the *car* package (Fox & Weisberg, 2018). To increase the likelihood of symmetrically distributed residuals, distribution of all variables was estimated visually using histograms, and data were transformed (default: natural logarithm, log_e_) when data distribution appeared asymmetrical. The covariate MFD was also log-transformed and both covariates of no interest, TIV and log_e_(MFD) were z-transformed to increase interpretability of the results (Schielzeth, 2010).

##### Post-hoc analysis of VBM results

Post-hoc analysis in significant VBM clusters was performed to confirm SPM analyses, in which the 2-by-2 design (group-by-time) was reduced to a two-sample t-test over the difference images (post-minus pre-intervention). Group-by-time interaction effects were tested in linear mixed models for each cluster separately, which allowed the investigation of regional differences and patterns. Since the effect of interest was the group-by-time interaction effect, the null model only included the main effects of group and time as fixed effects. The model equation is depicted below; β_1_…β_5_ denotes the two-way interaction and the main effects of all variables in interaction terms and the covariates, *u* and *e* depict random intercepts per subject and subject residuals. The GMV values from the individual clusters were log_e_-transformed. The *p* values from the full-null-model comparison were corrected using the *Holm-Bonferroni* method in the *p*.*adjust* function from the *stats* package.

Following a significant group-by-time interaction effect, post-hoc tests were conducted to test within-group effects using the *emmeans* function with *Holm-Bonferroni* correction from the *emmeans* package (Lenth, 2021).

Full model:

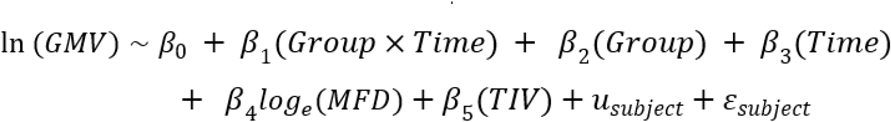

Null model:

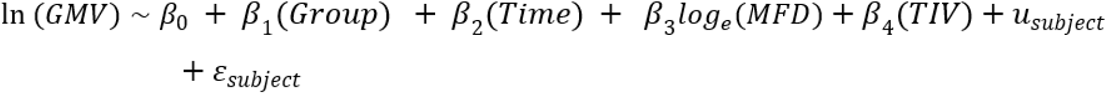

##### Total GM, total WM, and CSF

Only data of participants included in the VBM analysis were used.

Log_e_-transformation was applied to values of total GM and total WM, while total CSF was left untransformed (criterion: symmetry of the distribution of residuals; see above). The full models included the main effects and the group-by-time interaction (plus covariates log_e_(MFD) and TIV), while the null models lacked the interaction. Significant interaction effects were followed by post-hoc tests using the *emmeans* function.

##### Quantitative T1 values

Only data of participants included in the VBM analysis were used.

Before analysis, T1 values (mean per participants and cluster) were z-transformed and outliers of 3 SD above and below the sample mean were removed. Because 11 of the resulting 13 outliers came from the same two participants, these were excluded from the T1 analysis entirely (remaining sample: n = 61). T1 values were left untransformed (criterion: symmetry of the distribution of residuals; see above). The full model included the main effects, all two-way interactions, and the three-way interaction of group, time, and cluster (plus covariates log_e_(MFD) and TIV), while the null model lacked all interactions. If the three-way interaction was not significant, it was excluded from the full model (reduced model). The null model remained unchanged. Significant interaction effects were followed by post-hoc tests using the *emmeans* function.

##### T1-weighted intensity values

Only data of participants included in the VBM analysis were used.

Before analysis, T1-weighted intensity values of 3 SD above the sample mean were excluded as outliers. Since the resulting 6 outliers came from the same participant, he was excluded from the analysis (analysed sample: n = 62). The full model included the main effects, all two-way interactions, and the three-way interaction of group, time, and cluster (plus covariates log_e_(MFD) and TIV), while the null model lacked all interactions. T1-weighted intensity values followed a symmetrical distribution and were left untransformed.

##### Anatomical ROIs

We investigated group differences in GMV over time within the hypothesized four ROIs in four separate models (left and right amygdala, left and right hippocampus). As the effect-of-interest was the group-by-time interaction, the null model only included main effects of group and time. GMV values were log_e_-transformed to increase symmetry of variable distribution (see above).

#### CBF changes in the VBM clusters and in the whole brain

Following quality assessment, three participants were excluded from the pASL analysis. To investigate the impact of group and time on CBF within the VBM clusters, LMMs were set up in analogy to the VBM-ROI analysis. In addition to the factors group, time, cluster, and their interaction, a random effect per participant was included. The covariates TIV and MFD (included in the VBM-LMMs) were not included in the pASL analysis, because the preprocessing of the pASL data already included motion correction and TIV does not affect the intervention-induced change in CBF within a predefined region. (For a control analysis showing no significant effect of TIV, MFD or age on CBF data across all voxels from VBM clusters, see supplement, section 2.3.). CBF data followed a symmetrical distribution and was therefore not transformed before analysis.

As an exploratory analysis, group-specific CBF changes over time were assessed similarly to the VBM analysis, that is, groups were compared with a two-sample t-test on the difference images (post-pre) in SPM12. No nuisance variables were included, the sample-specific GM mask was used, and the threshold for TFCE correction was p_FWE_ < 0.05.

#### Analysis of endocrine, autonomic, and subjective stress measures

We investigated changes in autonomic (HR, HRV), endocrine (saliva cortisol, plasma ACTH), and subjective stress measures (STAI - state anxiety, VAS “stressfulness”) over time between groups using LMMs. All time points beginning from directly after the first (T3) until directly after the second (T11 and T12) structural MR scan were included (10 time points for endocrine and subjective data and 12 for autonomic data). One “non-responder” participant was excluded from the endocrine analysis due to a cortisol increase below 1.5 nmol/l (Miller et al., 2013), one data point was identified as a measurement error (value dropped by 98% to near 0 and then returned to 81%) by visual inspection of the residuals plot and excluded from the saliva cortisol model. The full model included group and time as well as their interaction and baseline (mean between 2 time points before intervention: T4 and T5 (-30min and -15min), see Figure 1) values as fixed effects and a random intercept per subject. Full models were compared against the respective null model lacking the interaction effect with X^2^ tests. Saliva cortisol, plasma ACTH, HR, and STAI score values were log_e_-transformed, HRV and VAS stressfulness values were square-root transformed. (More details on LMM analysis can be found in the section *Analysis of extracted imaging markers* above.)

#### Association of VBM changes with other stress measures

We conducted two types of analyses to investigate the association of GMV changes with endocrine, autonomic, and subjective stress measures: LMMs were used to analyse the effect of the trajectory of endocrine, autonomic, and subjective stress measures on GMV changes. Linear models (LMs) were used to test the association between stress reactivity and GMV changes by analysing the association of ΔGMV values (post-pre) and the peak reactivity value of the stress measures (maximum-baseline). Peak reactivity is commonly used in stress research (Engert et al., 2013; Van Cauter & Refetoff, 1985), also to determine individuals with a cortisol increase below physiological relevance (“non-responders”; Engert et al., 2013; Miller et al., 2013; Van Cauter & Refetoff, 1985).

LMMs have the advantage of covering the trajectory of stress measures by including data from all timepoints (before, during, and after the intervention). However, this high number of observations in stress measures also adds a lot of variance compared to GMV data, available at only two time points, which may overfit the model. Please note that both LMMs and LMs (with Δ values) are complementary analyses, which - since they are not built on the same data -cannot be directly compared with regard to variance explained (e.g., adjusted R^2^) or model fit (e.g., Akaike Information Criterion AIC).

P values from LMMs and LMs were multiple comparison-corrected using Holm’s method (Holm, 1979; 6 stress measures, 2 analyses, LMM and LM, each = 12) as implemented in the *p*.*adjust* function of the *stats* package. In case of significance, the *emtrends* function from the *emmeans* package (Lenth, 2021) was used to extract the within-group estimates and test for a significant interaction effect. The *drop1* function was used to extract the estimates and *p* values for the single predictors.

##### Association of VBM changes with other stress measures (LMMs)

Stress measures (saliva cortisol, plasma ACTH, HR, HRV, STAI score, and VAS score) were included in separate full-model LMMs and tested against a null model without them. The model equation is depicted below; denotes all two-way interactions and the main effects of all variables in the interaction term, *u* and *e* depict random intercepts per subject and subject residuals.

Full model:

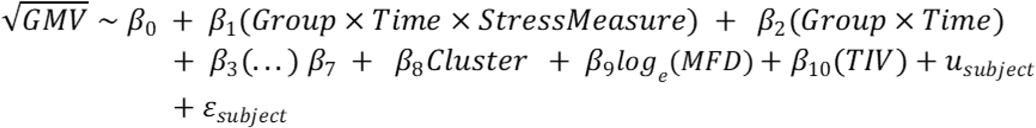

Null model:

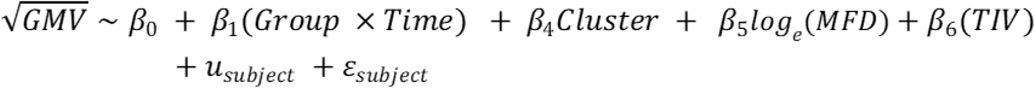

##### Association of ΔVBM with peak reactivity of other stress measures (LMs)

ΔGMV values were calculated by subtracting the pre-from the post-scan value. Δ values of stress-measures (saliva cortisol, plasma ACTH, HR, HRV, STAI score, and VAS score) were peak reactivity values calculated by subtracting the baseline value from the maximum value within 15-45 minutes after intervention onset (Engert et al., 2013; Van Cauter & Refetoff, 1985). The assumptions for LMs were tested as described above for LMMs.

ΔGMV was the dependent variable in all models. The full model included group and peak reactivity of stress measures as well as their interaction and the mean of pre- and post-TIV as well as MFD as independent variables. This was compared against a null model lacking the peak reactivity of stress measures using an F-test.

Full model:

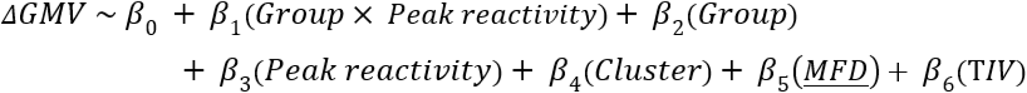

Null model:

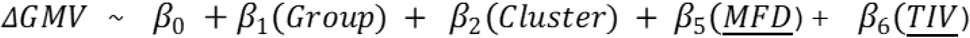

## Results

### Whole-brain VBM: significant interaction effect in 15 clusters

After quality assessment, the VBM was analysed in 63 participants: 30 in the stress group (TSST) and 33 in the control group (placebo-TSST). For an illustration of the final sample sizes for each parameter, see Figure S3. The results from the two-sample t-test on Δgrey matter images (ΔGM) were investigated using two-sided t-contrasts (i.e., control > stress and control < stress). The T contrast for control > stress did not yield statistically significant results. The opposite contrast (control < stress) showed a significant (p_FWE_ < 0.05) effect in 16 clusters (see Table 1, Table S1 and Figure 2), including cortical midline structures (CMS) and bilateral insula. (The cluster with an extent of 1 voxel was excluded from further analyses). The unthresholded result maps can be found at https://www.neurovault.org/collections/SFQXOIUB/.

**Table 1.** Results from the voxel-based morphometry (VBM) analysis on grey matter volume (GMV) in the VBM clusters. Depicted are hemisphere, cluster name (derived from CAT12’s “neuromorphometrics atlas”), cluster size in voxels, p_FWE_ after threshold-free cluster enhancement (TFCE) correction, coordinates in MNI space (x y z), and change pattern as identified by post-hoc tests (see Figure 2). The cluster with an extent of 1 voxel was excluded from further analyses. P < 0.05 indicates a significant group-by-time interaction effect. N = 63 (stress group: n = 30).

|  | Hemis-<br>phere | Cluster Name | Cluster<br>Size | p<sub>FWE</sub><br>(TFCE) | x y z | GMV Change<br>Pattern |
| --- | --- | --- | --- | --- | --- | --- |
| 1 | L | Superior Medial Frontal Gyrus | 2364 | 0.014 | -03 50 30 | Pattern 3 |
| 2 | R | (posterior) Insula | 2441 | 0.015 | 43 -12 04 | Pattern 2 |
| 3 | L | (anterior) Insula | 466 | 0.027 | -40 -08 06 | Pattern 3 |
| 4 | R | Angular Gyrus | 696 | 0.035 | 54 -63 28 | Pattern 1 |
| 5 | L | Parahippocampal Gyrus | 35 | 0.038 | 43 -12 04 | Pattern 3 |
| 6 | R | Inferior Occipital Gyrus | 118 | 0.042 | 52 -80 03 | Pattern 3 |
| 7 | L | Mid- Cingulate Cortex | 160 | 0.042 | -03 -24 46 | Pattern 1 |
| 8 | R | Cerebro- Motor- Area | 77 | 0.043 | -04 -06 57 | Pattern 1 |
| 9 | R | Lateral Orbital Gyrus | 79 | 0.045 | 36 39 -15 | Pattern 2 |
| 10 | R | Precuneus | 141 | 0.046 | 03 -50 57 | Pattern 1 |
| 11 | R | Frontal Pole | 52 | 0.046 | 24 63 03 | Pattern 3 |
| 12 | R | Superior Medial Frontal Gyrus | 44 | 0.047 | 06 52 04 | Pattern 3 |
| 13 | L | Superior Temporal Gyrus | 48 | 0.048 | -63 -10 04 | Pattern 2 |
| 14 |  | // | 1 | 0.048 | 51 -46 51 |  |
| 15 | R | Middle Frontal Gyrus | 22 | 0.048 | 45 46 12 | Pattern 3 |
| 16 | R | Middle Frontal Gyrus | 23 | 0.050 | 42 54 -02 | Pattern 3 |

**Figure 2.**
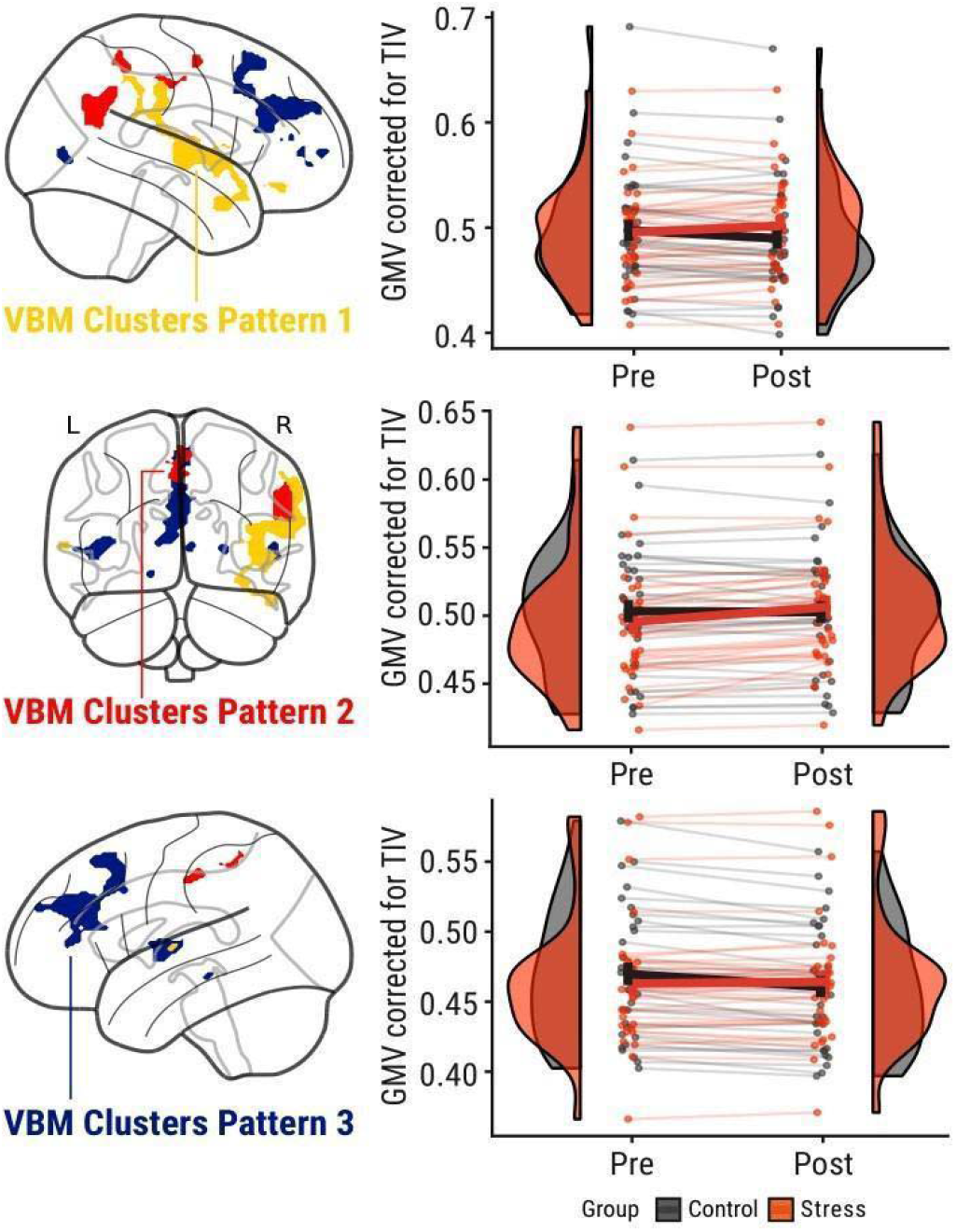
*Left column*: Voxel-based morphometry (VBM) results indicating a significant (p_FWE_ < 0.05) group-by-time interaction effect on grey matter volume (GMV). Colours indicate three distinguishable patterns: pattern 1 (yellow) - control: decrease, stress: increase; pattern 2 (red) - control: no significant change, stress: increase; pattern 3 (blue) - control: decrease, stress: no significant change. *Right column*: Changes in GMV group distributions (half violin) with individual changes (points, lines) and group means (central line with error bars). N = 63 (stress group: n = 30).

**Figure 3.**
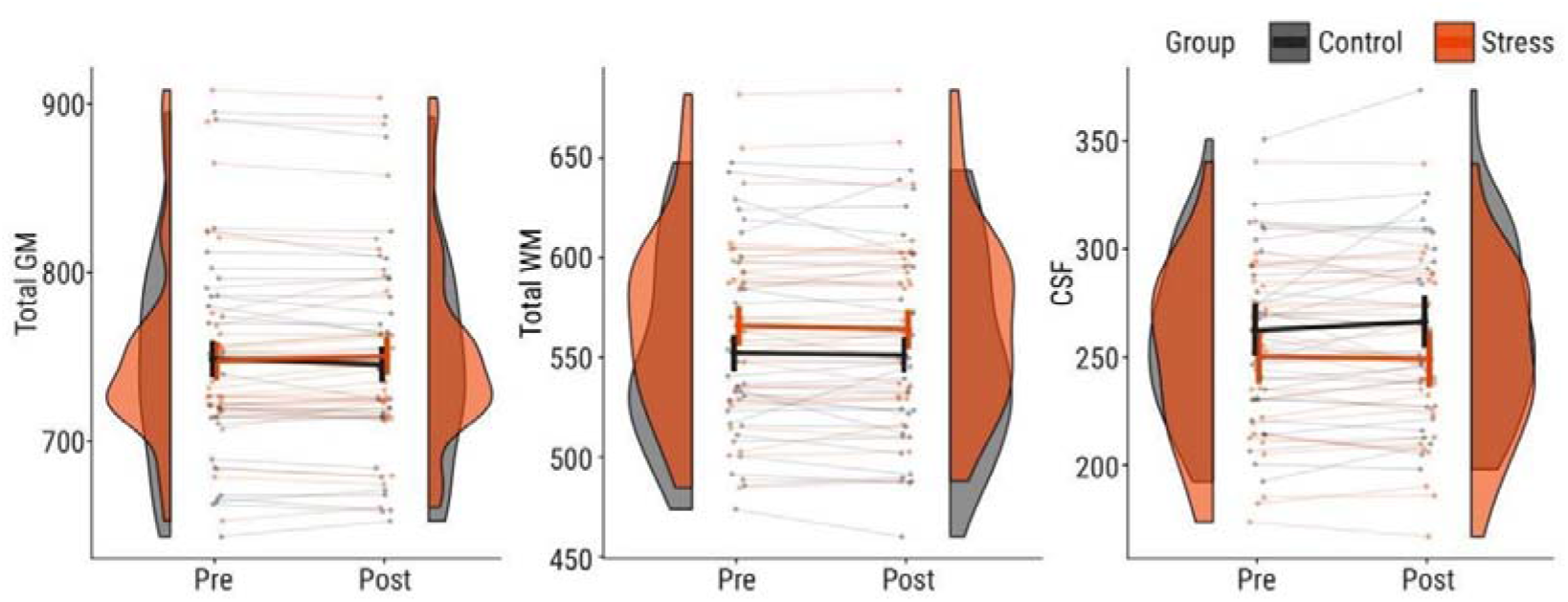
Change in total grey matter (GM), total white matter (WM), and total cerebrospinal fluid (CSF) volumes in the control group (grey) and in the stress group (red). Shown are scans (points) per subject (thin lines) and group distributions (half violin) for pre- and post-intervention scans. Bold lines indicate estimated marginal means and 95% confidence intervals obtained from linear mixed models. If data were transformed (log_e_) for statistical analysis, the estimates were back-transformed for visualization. N = 63 (stress group: n = 30).

### Post-hoc LMMs: three distinct change patterns

The VBM GM values from the whole-brain result clusters were extracted and the findings were tested in a post-hoc analysis using LMMs. In each individual VBM cluster, the full-null-model comparison showed a significant group-by-time interaction effect in all 15 clusters tested (see Table 1 for details). TIV explained a significant amount of variance (e.g., LSMFG: X^2^(1) = 23.38, p < 0.0001) in 13 clusters, while MFD did not (e.g., LSMFG: X^2^(1) = 0.60, p < 0.4381). Post-hoc tests revealed three patterns (Figure 2):

1. three clusters, including the Right Posterior Insula, showed a significant GMV increase in the stress group and a significant GMV decrease in the control group;
2. four clusters, including the Right Angular Gyrus and Left Mid-Cingulate Cortex, showed a significant GMV increase in the stress group and no significant change in the control group; and
3. eight clusters, including the biggest cluster in the anterior cortical midline (Left Superior Medial Frontal Gyrus) and the Left Anterior Insula, showed a significant GMV decrease in the control group and no significant change in the stress group.

### Significant group-differences in change of total GM and CSF volume

A significant group-by-time interaction effect was found for total GMV (Χ^2^(1) = 6.04, p = 0.0140) and CSF volume (CSFV; Χ^2^(1) = 4.7, p = 0.0305), while no significant change was found for WM volume (WMV; Χ^2^(1) = 0.20, p = 0.657). Post-hoc tests were not significant for total GMV (control: t/t_c_(60.3) = -1.96, p = 0.1084; stress: t/t_s_(62.9) = 1.70, p = 0.1084) but qualitatively showed a decrease in the control group (-0.4%, *β_c_* = -0.004) and an increase in the stress group (0.4%, *β_s_* = 0.004). CSFV increased significantly in the control group (1.5%, *β_c_* = 3.98, t/t_c_(60.2) = 2.6, p = 0.0232) and decreased non-significantly in the stress group (-0.4%, *β_c_* = −1.23, t/t_s_(62.3) = -0.7, p = 0.4857).

### Additional MR parameters in VBM clusters

#### GM T1 - but not T1w intensity values in GM increase in both groups

The group-by-time interaction effect found in the VBM data was not significant in the extracted T1 values (X^2^(1) = 0.405, p = 0.5246). There was a significant time-by-cluster interaction effect (X^2^(14) = 134.08, p < 0.0001), indicating an increase in T1 values over time in 7 of the 15 clusters, which did not include the three biggest clusters in the SMFG and bilateral insula (Table S4, Figure S7). On average, the T1 value increased (by ∼48 ms or ∼3.4%) with time across groups. As can be seen in Table S4, the average T1 value at the pre- and post-intervention scan in clusters with significant increases in T1 values (e.g., Right Angular Gyrus) was (1) lower than in clusters without significant increases in T1 values (e.g., Right Posterior Insula) and (2) lower than expected in GM at 3T (1350 ms, Marques et al., 2010), which may reflect a contribution of white matter (T1 at 3T: 810 ms; Marques et al., 2010) due to partial volume effects. For clarification, we conducted the following supplementary analyses (see supplement, section 2.2. for details): We tested whether extracted T1 values in GM voxels outside of the VBM clusters (but inside a GM mask) would also show a statistically significant change and found an increase of similar magnitude (by 29 ms or 2.4%; X^2^(1) = 20.50, p < 0.0001). A binary GM mask was created by combining all GM images from our sample and thresholding their values based on the probability of each voxel being located in GM (between 0 and 1). In the main analysis, a threshold of 0.1 (10% probability that the voxels are located in GM) was used, as recommended in the CAT12 manual (Version 30-06-2021, http://www.neuro.uni-jena.de/cat12/CAT12-Manual.pdf). We compared T1 values within GM masks at different thresholds (0.1, 0.2, 0.3, 0.5). A significant increase in T1 values was found at thresholds of 0.1, 0.2, and 0.3 but not at 0.5 (see Table S5), indicating that the T1 increase was driven by the edges of GM. That is, with an increasing threshold, the magnitude of the main effect of time decreased (0.1: 29 ms, 0.2: 24 ms, 0.3: 20 ms), and at 0.5 (2 ms), the main effect was no longer observed. Of note, when the mask with 0.5 GM probability was applied before extracting GMV values, the majority of clusters was masked out and only 4 clusters could be included in the analysis (LSMFG, bilateral Insula, and MCC), none of which had shown a significant increase in T1 values before thresholding. We also investigated whether the definition of GM boundaries (i.e., using GM masks with thresholds of 0.1, 0.2, 0.3, and 0.5) would affect our VBM results and found that 11 clusters were significant at a threshold of 0.3. The clusters in precuneus and the right posterior insula were robust even to a threshold of 0.5.

The group-by-time interaction effect found in the VBM data was not significant in the extracted T1-weighted intensity values (Χ^2^(1) = 3.13, p = 0.99). The main effect of time was not significant either (X^2^(1) = 2.16, p = 0.141, Figure 4), but a significant main effect of group (X^2^(1) = 6.9, p = 0.0087) indicated a difference in initial T1-weighted intensity values that remained constant over time (for follow-up analyses, see supplement, section 2.3).

**Figure 4:**
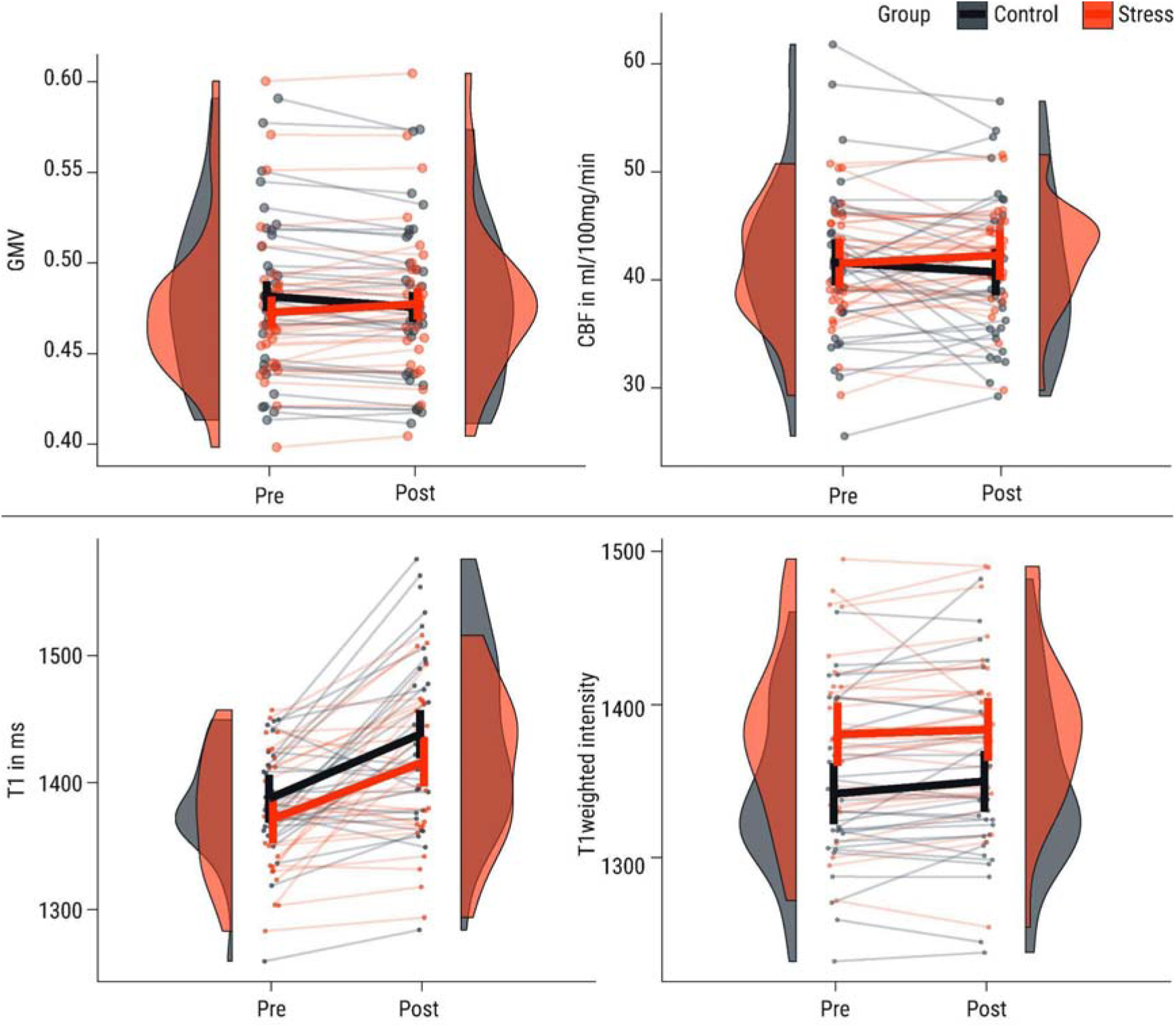
Change in grey matter volume (GMV), cerebral blood flow (CBF), T1, and T1-weighted intensity values in the control group (grey) and in the stress group (red). Shown are scans (points) per subject (thin lines) averaged across clusters and group distributions (half violin) for the pre- and post-intervention scan. Bold lines indicate estimated marginal means and 95% confidence intervals obtained from linear mixed models. If data were transformed (log_e_ or square-root) for statistical analysis, the estimates were back-transformed for visualization.

#### Cerebral blood flow is not significantly increased at 1 hour after stress

For CBF, there was a significant group-by-time interaction across all included clusters (Χ^2^(1) = 4.12, p = 0.0425). Post-hoc tests in both groups separately showed no significant effect in either group but indicated that CBF decreased in the control group (by 0.9 ml/100g/min or ∼2.2%, t/t_c_(942) = -1.57, p(cor) = 0.2332) and increased in the stress group (by 0.8 ml/100g/min or ∼1.93%, t/t_s_(942) = 1.296, p = 0.2332), resembling pattern 1 of the VBM results (Figure 4). However, the interaction was driven by two participants in the control group, who showed an exceptionally large decrease in CBF (Figure 4). When they were removed from the analysis, the interaction did not remain significant (Χ^2^(1) = 1.36, p = 0.2436) and CBF decreased only marginally in the control group (by 0.2 ml/100g/min or ∼0.4%, t/t_c_(912) = -0.324, p(cor) = 0.7463).

In cluster-specific post-hoc tests, no CBF changes survived multiple-comparison correction (all p_corr_ > 0.29). For more details of the CBF results, see supplement (Table S6 & S7, Figure S7).

### Amygdala and Hippocampus show no significant change in GMV

Comparing the full to the null model showed no significant group-by-time interaction effect on GMV in the left Amygdala (X^2^(1) = 0.60, p = 0.4372), right Amygdala (X^2^(1) = 0.77, p = 0.3803), left Hippocampus (X^2^(1) = 0.13, p = 0.7227), or right Hippocampus (X^2^(1) = 0.02, p = 0.8805).

### Robust stress response in autonomic, endocrine, and subjective stress measures

The TSST induced a robust stress response in autonomic, endocrine, and subjective stress measures, as also shown in previous publications from our study (Reinelt & Uhlig et al., 2019 and Bae & Reinelt et al., 2019). Significant (p_corr_ < 0.05 with Bonferroni-Holm correction) group-by-time interaction effects were present in all investigated autonomic, endocrine, and subjective markers (Table 2 and Figure 5).

**Table 2.**
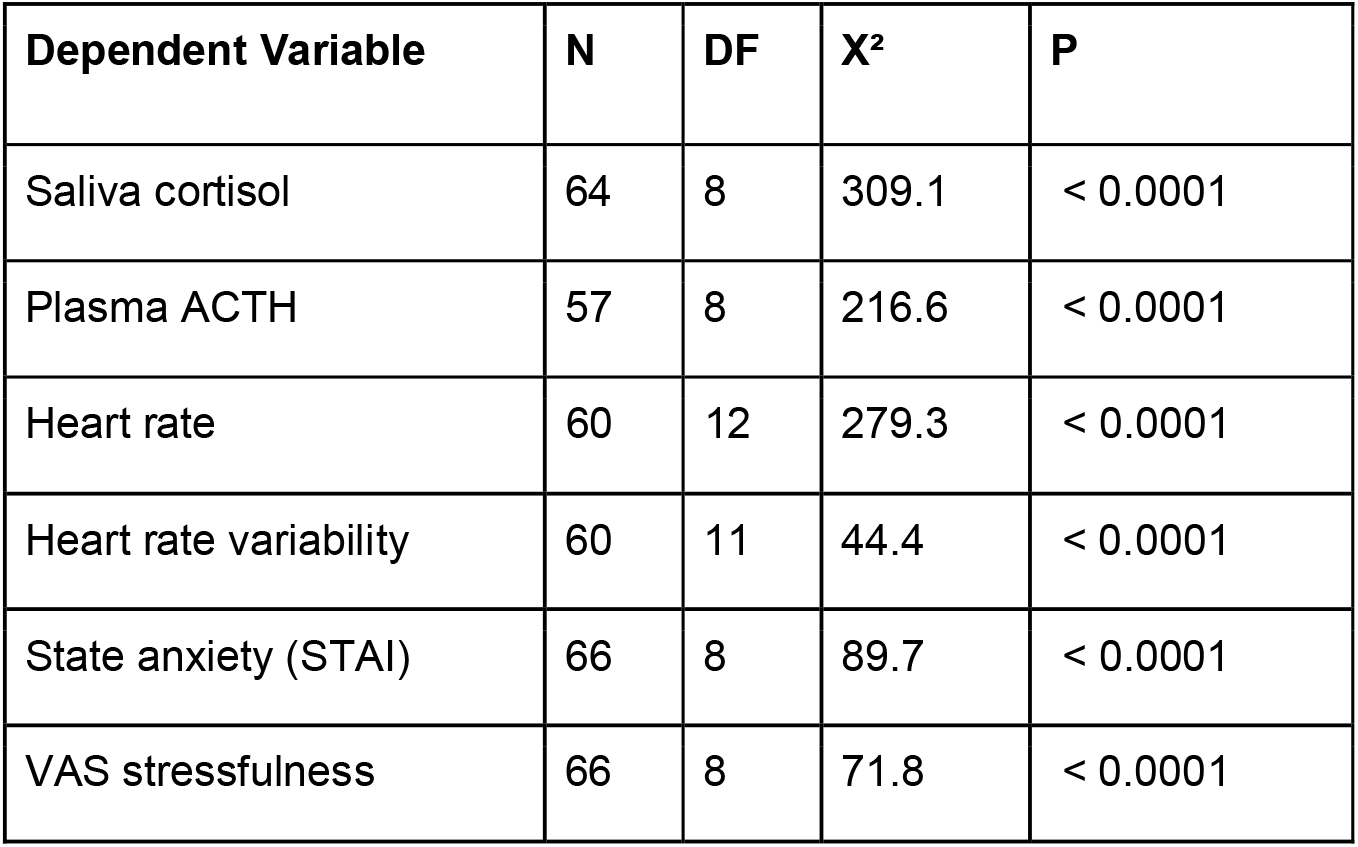
Results from linear mixed models on autonomic, endocrine, and subjective stress measures. Statistical parameters were obtained from a full-null-model comparison. Fixed effects: time, group, group-by-time interaction (full model only); Random effects: participant. Depicted are degrees of freedom (DF), the X^2^ and the *p* value from the full-null-model comparison. ACTH = adrenocorticotropic hormone; STAI = State Anxiety Inventory.

**Figure 5:**
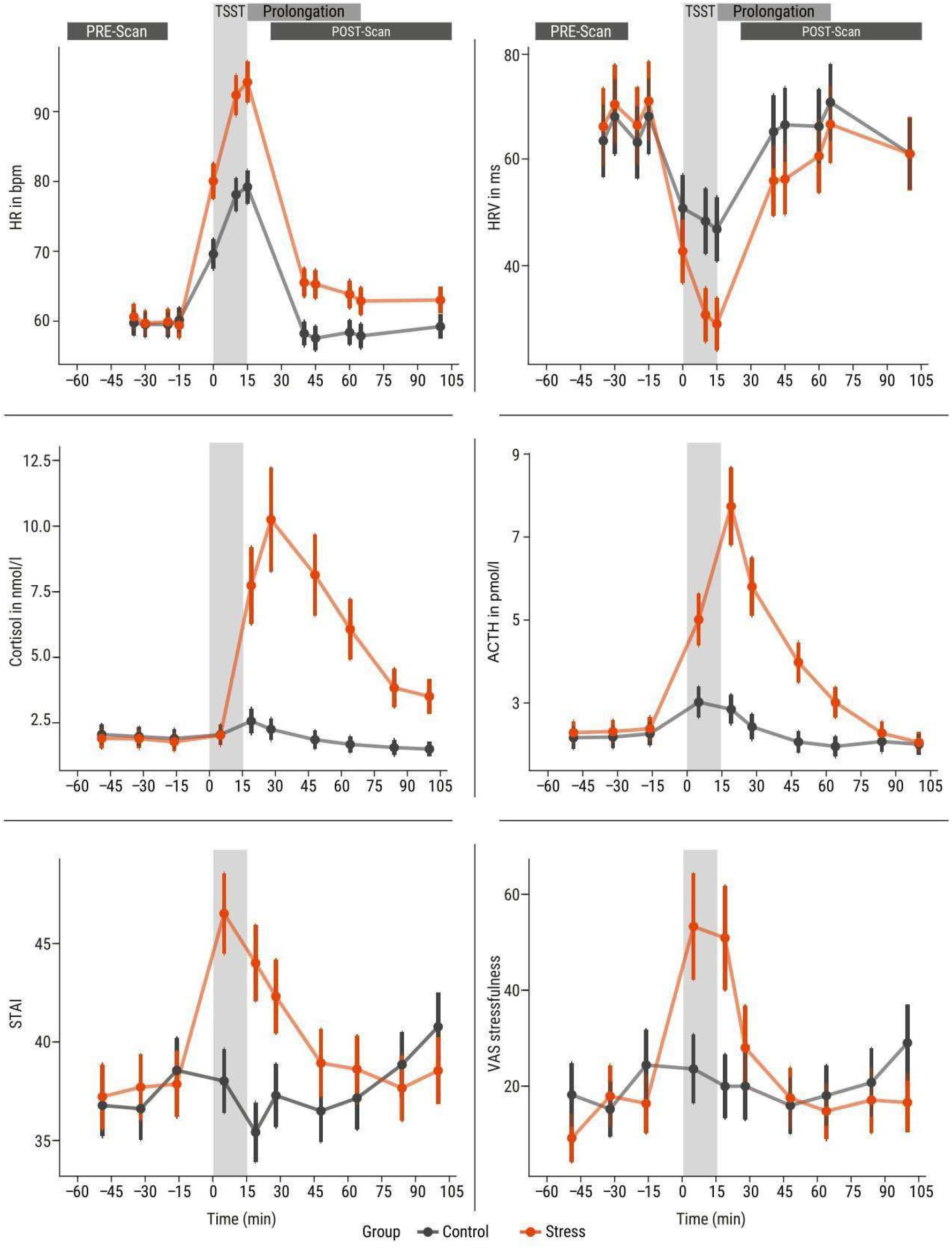
Time courses (x-axis: time of day) of saliva cortisol (nmol/l) and plasma adrenocorticotropic hormone (ACTH) (pmol/l) concentrations, heart rate (beats per minute) and heart rate variability (RMSSD in ms) and subjective stress measured by state anxiety (State Anxiety Inventory, STAI, sum score) and a visual analogue scale (VAS, score) of stressfulness. Plotted are the estimated marginal means from the linear mixed models (see above). If data were transformed (log_e_ or square-root) for statistical analysis, the estimates were back-transformed for visualization. Error bars depict upper and lower 95% confidence intervals for model estimates. Grey: control group, orange: stress group. (Only timepoints between the two structural scans are included; for the full time courses and their statistical analysis, see Bae & Reinelt, et al., 2019; Reinelt & Uhlig, et al., 2019.)

Post-hoc tests and visualization (Figure 5) show the dynamics of the stress response: subjective stress peaked earliest (+5 min) and saliva cortisol latest (+25 min). Heart rate was the first parameter to return to baseline (+25 min) while the group difference in saliva cortisol remained longest (+90 min).

### Association of GMV with other stress measures in VBM clusters

#### LMM: no significant association of VBM changes with other stress measures

After multiple-comparison correction, no significant association of GMV changes with autonomic (HR and HRV), endocrine (saliva cortisol and plasma ACTH), and subjective (STAI score and VAS score) stress measures was found in any cluster in the LMM analysis (see Supplementary Table S3 and Figure S6 for details).

#### LM: ΔGMV is significantly correlated with HRV and STAI peak reactivity

##### Endocrine stress measures

In the full-null-model comparison, there was no significant effect of saliva cortisol (F(2) = 0.63,p_corr_ = 1) or plasma ACTH peak reactivity (F(2) = 2.03, p_corr_ = 0.1321) on ΔGMV (Figure S6).

##### Autonomic stress measures

In the full-null-model comparison, there was no significant effect of HR peak reactivity (F(2) = 3.51, p_corr_ = 0.2736, Figure S6) on ΔGMV. HRV peak reactivity (F(2) = 6.24, p_corr_ = 0.0224) was significantly associated with ΔGMV. Post-hoc tests showed no significant interaction effect for group (t/t(827) = -1.59, p = 0.1107), but a negative association between HRV peak reactivity and ΔGMV both in the stress (*β_s_* = -0.0000327) and the control group (*β*_*c*_ = -0.0000885). In both groups, the participants who showed more pronounced HRV decreases also showed stronger GMV increases (or weaker GMV decreases, Figure 6).

**Figure 6:**
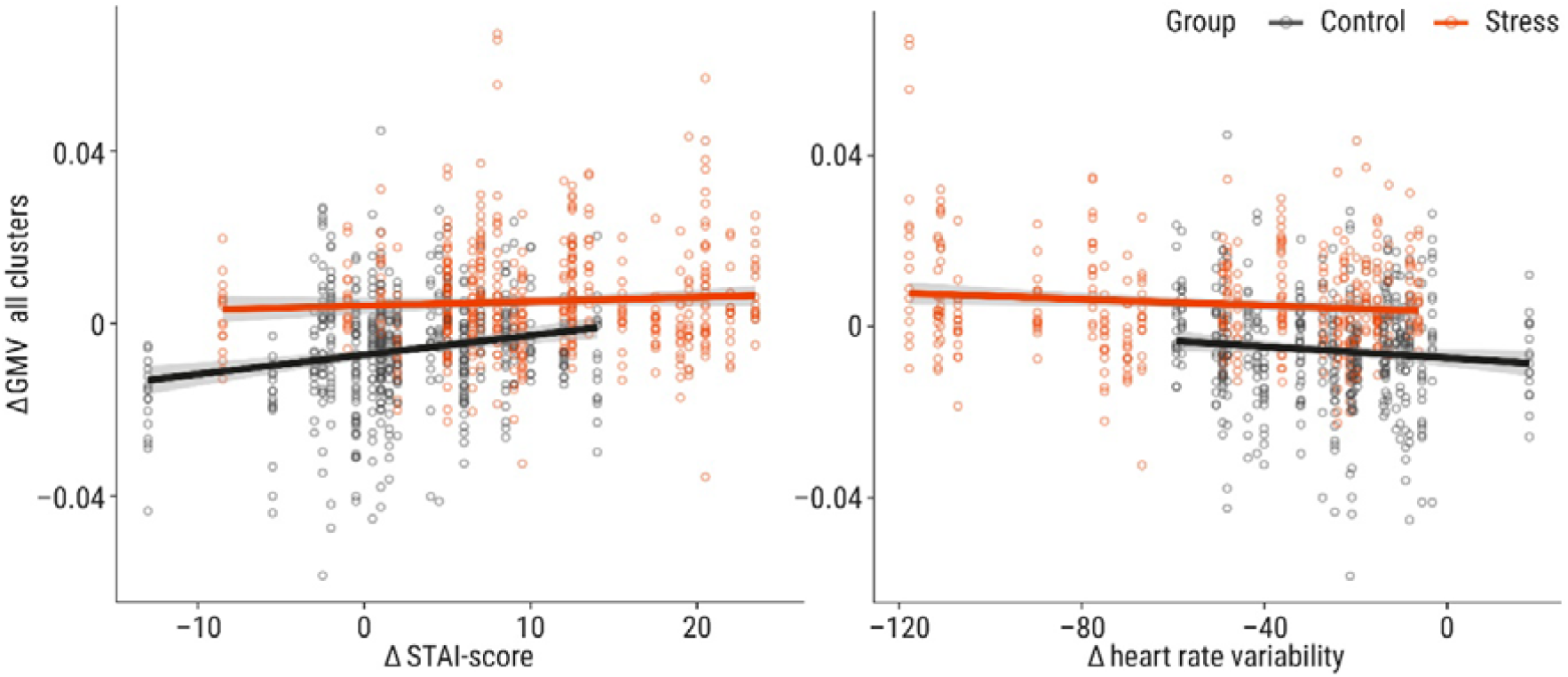
Association of Δgrey matter volume (GMV; post-pre) with peak reactivity of stress measures. Shown are significant associations from linear models (LMs): state anxiety (State Anxiety Inventory, STAI; positive association) and heart rate variability (RMSSD; negative association). The LMs revealed no significant association with saliva cortisol, ACTH (Adrenocorticotropic hormone), heart rate, and subjective stressfulness (Table S3, Figure S6). Line indicates slope and standard error. Points indicate GMV values per voxel-based morphometry (VBM) cluster and subject, each subject is represented in one column of points. Grey: control group, orange: stress group.

##### Subjective stress measures

In the full-null-model comparison, there was a significant effect of STAI score peak reactivity on ΔGMV (F(2) = 7.586, p_corr_ = 0.0065). Post-hoc tests showed a significant interaction effect with group (t/t(987) = 2.335, p = 0.0197) and a positive association between STAI score peak reactivity and ΔGMV in the stress (*β*_*s*_ = 0.0000902) and – even stronger – the control group (*β*_*c*_ = 0.0003895). In both groups, the participants who showed more pronounced STAI score increases also showed stronger GMV increases (or weaker GMV decreases, Figure 6).

In the full-null-model comparison, there was no significant effect of VAS stressfulness peak reactivity of ΔGMV (F(2) = 3.879, p_corr_ = 0.1894, Table S3, Figure S6).

## Discussion

Using voxel-based morphometry, we found rapid volumetric brain changes that differed between groups over time in 15 clusters, mainly along the cortical midline and in the bilateral insula. We identified three patterns of GMV changes across the clusters: the stress group showed a GMV increase (patterns 1 and 2) or no change (pattern 3) while the control group showed a GMV decrease (patterns 1 and 3) or no change (pattern 2). Our stress intervention induced a pronounced stress response on the autonomic, endocrine, and subjective levels. Changes in GMV were related to peak reactivity in state anxiety and heart rate variability but not in heart rate, saliva cortisol, plasma ACTH, or subjective stressfulness.

To explore the microstructural and physiological basis of these findings, we also analysed quantitative T1 and CBF imaging parameters. The significant group difference over time was not present in T1 or T1-weighted intensity values. In T1 values, a significant increase over time across groups occurred, which post-hoc tests showed to be significant in half of the clusters. CBF across VBM clusters non-significantly decreased in the control group and increased in the stress group. Thus, the stress-related brain changes are reflected in local GMV increases relative to the control group. In clusters with no significant GMV change, the increase may be masked by the GMV decrease observed in the control group. We did not observe the hypothesized GMV changes in hippocampus and amygdala.

In summary, we found that the dynamics of rapid volumetric brain changes differed between groups, suggesting that endogenous brain changes (GMV decrease) are counteracted by acute stress.

The rapidness of brain changes we detected with structural MR imaging methods raises the question of their physiological origins. Mouse studies have connected VBM changes to altered dendritic spine density: Aversive, stressful stimulation, like auditory fear conditioning (Keifer et al., 2015) or restraint (Kassem et al., 2013) led to volumetric changes, measured with volumetric MRI (Kassem et al., 2013) or VBM (Keifer et al., 2015), which were correlated with spike density changes in functionally relevant regions, such as amygdala and insula (Keifer et al., 2015) as well as ACC (Kassem et al., 2013). Synaptic and dendritic plasticity may be detectable after minutes to hours (Johansen-Berg et al., 2012); however, we would expect subtle T1 shortening from such processes due to an increase in the amount of membranes and macromolecules and concomitantly reduced water content (Fullerton et al., 1982) whereas longer T1 values were observed in both groups with masks thresholded at GM probabilities ≤0.3. Changes to dendritic morphology may further be accompanied by migration or swelling of capillaries and glia in order to compensate for heightened energy demand resulting in increased tissue volume, which manifests itself as GMV changes detected by VBM (Lövdén et al., 2013). Glial cells have been shown to react to sensory stimulation (Tremblay et al., 2010) and can modulate plasticity and learning processes (Jammal et al., 2018).

Especially astrocytic glia cells are prominent candidates for targeting cellular structures in brain plasticity. These non-myelinating glia cells are involved in neuronal metabolism and fluid homeostasis, and they can mediate the excitability of neurons (Shao & McCarthy, 1994). Activation may cause astrocytes to swell within seconds or minutes, which has been shown to affect diffusion-weighted MRI measures (Johansen-Berg et al., 2012) and may also affect estimates of GMV. Astrocytes also express corticosteroid receptors (Bohn et al., 1991), and their structure and function can be influenced by chronic (Tynan et al., 2013) as well as acute stress (Braun et al., 2009). Stress-induced astrocyte plasticity has also been linked to stress-related psychiatric diseases (for reviews, see Bender et al., 2016; Cathomas et al., 2022).

Thus, the observed GMV changes may reflect (transient) local tissue changes and/or vascular changes to accommodate changes in energy demand following neural activity. These alterations, which are present more than an hour after the stress episode, may also be related to the induction of – potentially long-term – morphological changes.

In addition to a stress-induced increase in GMV, we also found local GMV decreases in the control group. A linear decrease in total GMV of similar magnitude (∼1%) from morning to afternoon has been shown in Karch et al., 2019 and Trefler et al., 2016, which was accompanied by regional GMV changes, for example, in the MPFC and precuneus (Trefler et al., 2016). In our study, the two structural scans were separated by approximately 2.5 hours, from early to late afternoon. We also found total GMV to decrease and CSF to increase from early to late afternoon in the control group, following the pattern of circadian rhythm-related GMV changes reported in Trefler et al. (2016). In the stress group, total GM and CSF non-significantly in the opposite direction compared to the control group: GM increased and CSF decreased. Exogeneous behavioural interventions have been shown to attenuate endogenous daytime effects on GMV (Trefler et al., 2016; Thomas et al., 2016) and so may a stressful intervention like the TSST.

We thus speculate that processes related to the circadian rhythm (i.e., supporting diurnal brain homeostasis; Trefler et al. 2016) contribute to the changes in our control group, while in the stress group, the behavioural intervention counteracts these processes. However, since we chose an active control group, we cannot rule out the possibility that the control intervention triggered brain changes detectable with VBM.

Extracted T1 values (at GM > 0.1) showed a significant increase over time in both groups and in seven of the 15 VBM clusters but no significant group difference, that is, the VBM interaction was not mirrored in T1 values. Moreover, diminishing T1 increases were obtained with increasing GM thresholds until no changes remained at a threshold of 0.5. This suggests a stronger contribution of CSF through partial volume effects in those clusters in the post-compared to the pre-scan. This “apparent” increase in T1 values (thresholded at GM > 0.1) was not limited to the VBM clusters but occurred in all GM, probably driven by effects at the GM boundaries. Decreased GMV along with increased CSFV has been reported in association with daytime (Trefler et al., 2016) but also following dehydration (Streitbürger et al., 2012). We minimized the variability of food and fluid intake by providing a standardized lunch, but we did not measure the participants’ hydration status and cannot exclude the possibility of group differences in hydration. Yet, in Streitbürger et al. (2012), dehydration mainly affected GMV in areas close to the ventricles rather than cingulate and insular cortices (the main VBM clusters in our results), and dehydration-induced effects should occur in both groups alike. However, while T1 increased in both groups, a significant increase in total CSFV was found in the control group but not in the stress group. It is possible that differentially altered cellular volume following water migration is reflected in an increase in T1 as well as in a differential change in tissue volume estimation by VBM. It has previously been shown that neural activity is associated with cell swelling and accompanied by a fluid shift from extra- to intracellular space (Sykova 1997), and it has been proposed that such processes could be picked up by VBM as an apparent increase in GMV (Naegel et al, 2017).

Applying our assumption about an endogenous homeostatic process being counteracted by processes accompanying stress-induced neural activity here, we can speculate about the following mechanisms: 1) endogenous fluid shifts, which occur in both groups (possibly hydration- or daytime-related), are reflected in an apparent increase in T1 values. These changes are accompanied by 2) cell shrinkage in the control group, reflected in a VBM-estimated decrease in GMV and increase in CSF volume (CSFV) as well as 3) cell swelling (following stress-induced neural activity) in the stress group, reflected in a VBM-estimated increase in GMV and decrease in CSFV, with simultaneously increased T1 values. Different patterns of fluid shifts between compartments may thus explain the divergence between the T1 and the VBM results. It needs to be noted that such inter-compartmental fluid shifts remain speculative as they happen too quickly to be picked up by our MRI sequences.

In summary, GMV decreases in the control group may reflect changes in fluid homeostasis (e.g., associated with the circadian rhythm) along with cell swelling or shrinkage. In the stress group, such processes may be counteracted by processes that regulate the energy demand following neuronal activation in response to the stressful intervention. This increased energy demand following brain activity under stress may also be reflected in increased CBF, which has been shown to affect VBM measures of GMV (Ge et al., 2017).

Consistent with the overall GMV decrease in the control group, we find non-significantly decreased CBF in that group across all clusters (although this appears to be mainly driven by few participants). We also found CBF increases in the stress group in the left and right SMFG, which, however, also did not survive multiple-comparison correction. CBF increases have previously been shown (using ASL) during an in-scanner stressor, for example in the right PFC, ACC, insula, and putamen (Wang et al., 2005). Many brain vessels are located along the medial wall, including the middle cerebral artery, but also the insula displays a particularly high density of vessels, including the anterior cerebral artery (Mouches & Forkert, 2019). Thus, our main VBM clusters (bilateral SMFG and insula) are in the vicinity of major vessels. During stress-induced physiological activity, changes in blood parameters (e.g., blood flow) and vasodilation could influence the VBM analysis. However, we did not find significant stress-induced CBF increases. One reason may be the limited sensitivity of the pASL analysis due to the relatively low resolution, the time delay to the intervention (∼90 min) or limited spatial coverage (Figure S2). On the other hand, previous studies have shown overlapping but incongruent patterns of CBF and VBM changes (Ge et al., 2017, Franklin et al., 2013), which may indicate that other processes, such as changes in brain metabolites in response to functional activation, may affect the T1-weighted signal and thus contribute to apparent GMV changes measured with VBM (Ge et al., 2017). Especially in highly vascularized areas, hemodynamically induced GMV changes may also arise from changes in cerebral blood volume, (Kim and Ogawa, 2012), which we did not assess.

In addition, it has been proposed that an intervention-induced increase in oxygen-demand in specific brain areas may similarly affect estimations of GMV and decrease T1 values through changes in CBF and tissue oxygenation (Tardif et al., 2017). Here, we find no intervention-specific effect but increased T1 values in both groups. Given the time delay of > 1hr to the intervention, hemodynamic changes may have normalized until the MP2RAGE and pASL scan.

In summary, while changes in CBF mirror the pattern of our VBM results, we find limited evidence for a hemodynamic origin of GMV changes. Previous studies have found incongruent patterns of BOLD activation and volume or thickness changes (Olivo et al., 2022, Zaretskaya et al., 2022).

Functionally, the main clusters of stress related VBM changes in cortical midline structures (CMS) and bilateral insula can be related to the processing of emotional/stressful and self-relevant information.

The biggest cluster extended from the superior medial frontal cortex to the anterior cingulate cortex. Functionally, the medial frontal cortex has been involved in emotion processing (Etkin et al., 2011) and in the regulation of the physiological and behavioural stress response (McKlveen et al., 2015). It also has a high density of glucocorticoid receptors, which are central to the negative feedback mechanism of the HPA axis (Buchanan et al., 2010). Yet, we found no significant association of GMV with endocrine stress measures.

A significant association was found with the subjective and autonomic stress measures of state anxiety (STAI) and HRV, respectively. Decreased HRV correlated inversely with GMV changes in both groups, suggesting that control participants whose parasympathetic activity changed similarly to participants in the stress group showed less decrease in GMV than other control participants. In parallel, higher state anxiety was associated with less GMV decrease in both groups, but even stronger in the control group. These results indicate that parasympathetic deactivation and state anxiety, which can result from psychological stress, are linearly associated with GMV changes, and counteract the GMV decrease. In general, CMS – especially anterior ones – have been associated with self-relatedness and self-relevance (for a review, see Northoff & Bermpohl, 2004), a feature of any stressor and particularly of the TSST, in which participants “apply” for their individual dream jobs. Although participants knew the job interview was not real, they showed pronounced stress responses. Negative self-relevant stimuli and psychosocial stress have been shown to increase activity in CMS (e.g., MPFC; Lemogne et al., 2011) as well as connectivity between the amygdala and CMS (Veer et al., 2011), respectively.

In our study, two major clusters of stress-related GMV changes showed peaks in the left anterior insula and the right posterior insula. The insula has been understood as primary viscerosensory or interoceptive cortex with a posterior-to-anterior gradient (Craig, 2002): pain, temperature, and other homeostatically relevant bodily stimuli enter the posterior insula before they are integrated with other (e.g., exteroceptive) information and evaluated in the anterior insula, influencing subjective experience and guiding behaviour (Craig, 2002). The insula is also highly connected and often co-active with frontal CMS (e.g., MPFC and ACC), where the strongest cluster of stress-related GMV changes was found in our study, and they constitute a central axis of the salience network, which processes homeostatically relevant stimuli (Seeley, 2019). The integrative, multisensory function of the insula is also supported by animal studies showing, for example, that the posterior insula can shift behavioural strategies upon the detection of aversive or stressful interoceptive states (Gehrlach et al., 2019).

We have previously shown increased connectivity in the thalamus in response to stress (Reinelt & Uhlig et al., 2019), also to medial frontal regions and the insula (exploratory seed-based connectivity analysis in Reinelt & Uhlig, 2019), which showed stress-related GMV changes in the present study. The thalamus and the insula are part of the salience network (Hermans et al., 2014) and have been linked to interoception (thalamus: Barson, Mack, & Gao, 2020; Dobrushina et al., 2021; insula: Craig, 2002) as well as autonomic nervous system regulation (thalamus: Buijs, 2013; insula: Thayer & Lane, 2000). We can thus speculate that the increase in thalamic centrality reflects its role as a central hub for resource allocation (Garrett et al., 2018) to a variety of regions, some of which also show GMV changes. As an exploratory follow-up analysis, we investigated GMV changes in the thalamus cluster derived from the significant group-by-time interaction effect on EC values (Reinelt et al., 2019). This revealed no significant interaction effect, but a decrease in GMV across both groups in the thalamic cluster (For details, see supplement, section 2.1.2.).

In contrast to our hypothesis, we did not find significant GMV changes in hippocampus and amygdala. In the framework by Hermans et al. (2014), during and after acute stress, resources are allocated to the salience and (estimated at 1 hour after stressor onset) the executive control network, respectively. The post-intervention MRI was acquired 90 mins after stressor onset, which may coincide with the “downregulation” of the salience network (Hermans et al., 2014). Other studies have suggested an even earlier deactivation of limbic structures including the hippocampus and the amygdala during stress exposure (Pruessner et al., 2008)

In animal models of acute stress (Kassem et al., 2013, Chakraborty et al. 2020) and in stress-related mental disorders in humans (Chen et al., 2006; Karl et al., 2006) stress-induced brain structural changes in hippocampus and amygdala are found. At subclinical levels of chronic stress however, some studies did find changes in GMV (Dedovic et al., 2010; Savic, 2015; Suffren et al., 2021; Spalletta et al., 2014), but others did not: GMV reductions associated with stressful life events were, for example, found in the ACC, hippocampus, and parahippocampal gyrus, but not in the amygdala (Papagni et al., 2011) as well as in the MPFC and right insula, but not in the hippocampus or amygdala (Ansell et al., 2012). Possibly, GMV alterations in hippocampus and amygdala may be related to pathophysiological processes in the context of chronic or severe stress (Ansell et al., 2012) rather than the brain response to acute stress.

### Limitations

There are several limitations to our study. VBM can be considered a physiologically coarse method, and, despite several candidate processes (discussed above), the physiological origin of GMV changes remains unclear. VBM has also been criticized for introducing bias and neglecting non-linear effects, which are more pronounced when comparing heterogeneous groups (Bookstein, 2001; Davatzikos, 2004). Another limiting factor of our analysis is the spatial resolution. While a smoothing kernel of 6-8mm is recommended (see CAT12 manual) to optimize the data distribution and reduce the number of comparisons in a whole brain analysis, this reduces spatial accuracy. To estimate the influence of the smoothing kernel size, we repeated the VBM analysis with a smaller smoothing kernel: Decreasing the smoothing kernel to 6mm isotropic (FWHM) results in overall fewer and smaller clusters within the same main result regions as with the 8-mm smoothing kernel (see supplement, section 2.1.1.2.). By comparing two (randomly assigned) groups from a homogenous sample in our study, we expected to minimize such potential biases. The inclusion of young, healthy, male participants allowed us to investigate stress-induced changes using a multimodal approach without confounds like the impact of the ovarian cycle. However, the generalisability of our results remains to be tested in studies with more heterogeneous samples. The significant group-by-time interaction effect on GMV suggests that the differences are intervention-induced. While we kept the procedure as similar as possible between groups, we extended the TSST stressor by telling participants in the stress but not the control group there would be another task. Thus, it is possible that not the TSST alone but the prolongation of the stressor (or stress-related vigilance) in the stress group accounts for the group difference in GMV. Furthermore, a higher temporal resolution would add information about the trajectory of changes and about possible immediate transient changes and the stability of changes we observed in the stress and in the control group. We asked participants to refrain from drinking coffee in the morning to avoid caffeine effects on HPA axis activity (Patz et al., 2005). However, coffee can be an important part of the morning routine and its absence might have psychological (e.g., well-being) and physiological (e.g., metabolism) effects on regular coffee drinkers. Since we did not acquire information on coffee consumption habits, we cannot quantify such effects as well as potential group differences. We specifically investigated the effects of psychosocial stress in this study. To what extent findings generalize to other stressor types (e.g., physical stress) remains unclear. Head motion is a major neuroimaging confound (Beyer et al., 2020.), and it can decrease measures of GMV (Reuter et al., 2015). We aimed to physically minimize head motion during data acquisition and included the realignment / motion parameter MFD from the preceding resting-state scans as a proxy covariate in the VBM analyses. Head motion parameters (e.g., using gyrometry or video-based measures) from the actual MP2RAGE scan could be acquired using additional hardware.

## Conclusion

We find rapid brain changes following a psychosocial stress intervention compared to a placebo version of that task. Brain changes are observed in areas associated with the processing of emotional and self-relevant information but also with regulating HPA axis activity and sympathetic arousal. Stressed participants additionally show (non-significantly) increased cerebral blood flow in prefrontal areas. While CBF mirrors the VBM changes, neither T1, T1-weighted intensity, nor CBF fully account for the observed group differences over time in GMV. Our findings of rapid GMV changes following acute psychosocial stress detected with MRI in humans emphasize the influence of stress on the brain, suggesting that diurnal mechanisms of brain homeostasis are perturbed by acute stress.

## Acknowledgements

The visualization functions for LMM diagnostics were kindly provided by Dr. Roger Mundry.

## Data and code availability statement

The data that support the findings of this study are openly available at https://osf.io/vjyan/. In agreement with participant consent, this includes derived data, which cannot be used to identify individual participants. The code to reproduce the analyses can be found at https://gitlab.gwdg.de/necos/vbm.git.

## Footnotes

1 While we use the term “grey matter volume” for VBM changes, we consider it a placeholder, as other physiological changes may contribute to the signal (see below).

