## Supplementary Material for "Rapid volumetric brain changes after acute psychosocial stress"

### Supplement

#### 1. Methods

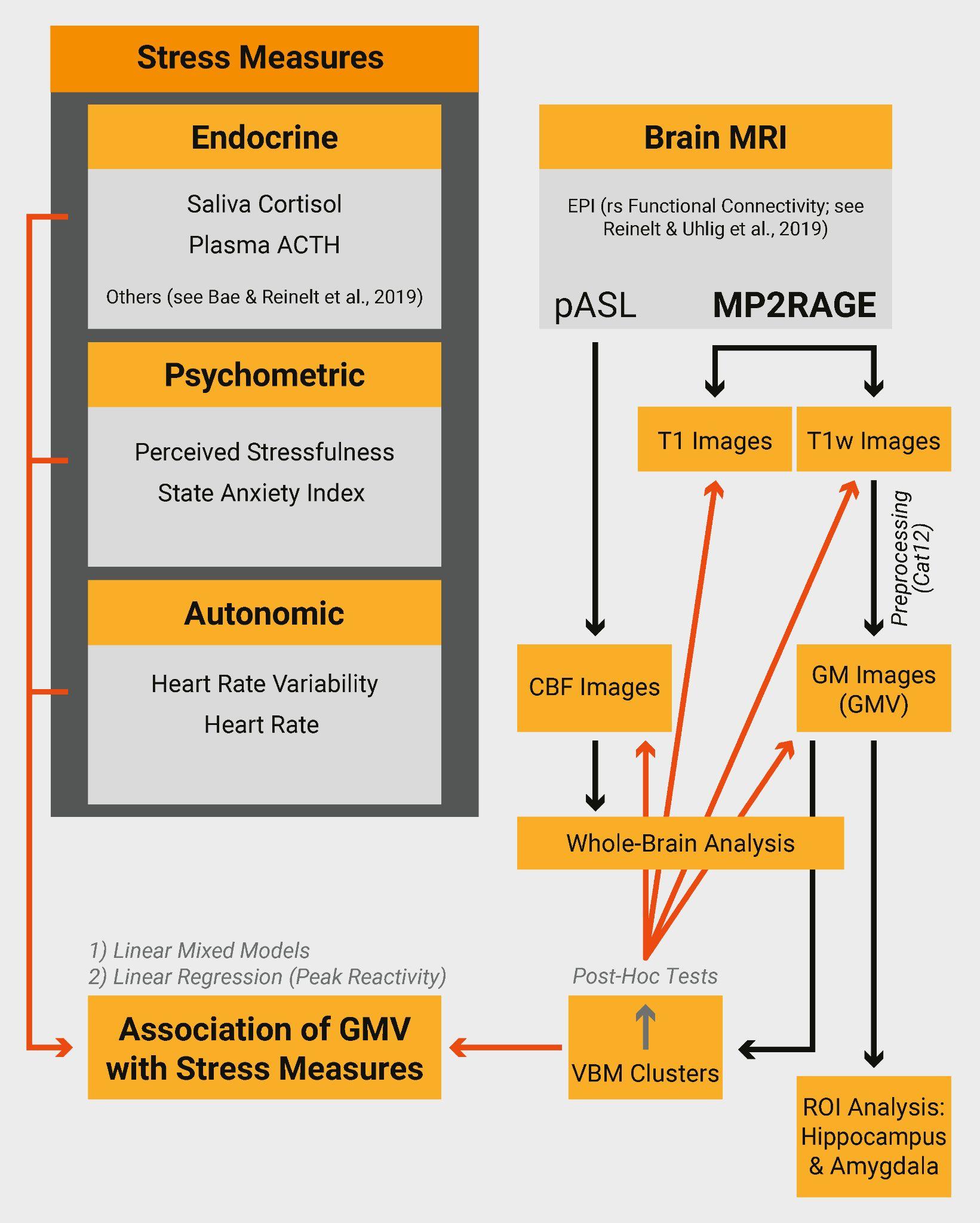

**Figure S1: Overview of analysis pipeline.** Black arrows = derived data, red arrows = post-hoc tests. Abbreviations: GMV =Grey Matter Volume, ROI= Region of Interest, CBF= Cerebral Blood Flow

##### 1.1. TSST

In a separate testing room, participants encountered a committee of two professional actors (one female, one male) introduced as two professional psychologists trained in the analysis of nonverbal communication. Participants were told that they should apply for their dream job (the job type had been inquired before) by freely describing their personal qualifications. While talking, they would be observed by the committee and recorded by a video camera and microphone. Furthermore, participants were told that they would have to perform another task after their talk and that they would have 5 minutes time to prepare notes for their oral introduction. Participants then sat down in front of the committee and could take notes for 5 minutes.

After this preparation phase, a sampling time point followed (T6), before the participants were instructed to stand in front of the microphone and to begin with their self presentation without using their notes. The committee members monitored the participant neutrally throughout the speech without any facial cues. Whenever the instructions were violated, committee members interrupted and repeated standardized instructions: for example, whenever the participants stopped talking, the committee members waited for 20 seconds, silently looking at the participants before asking them to resume their presentation. After 5 minutes, the participants were introduced to the next task, in which they had to count backwards in steps of 17 from 2043 as fast and accurately as possible. Every time they made a mistake, they had to start from the beginning. After 5 minutes, participants were told that they would be brought back to the MRI area, where another task would follow. Before that, they sat down for another sampling time point (T7). Afterwards, the participant was accompanied by the experimenter and the committee members to the MRI area. The procedure was chosen to maintain the subjective experience of stress and uncontrollability. After the second post-stress resting-state measurement (i.e., before the post-stress MP2RAGE), participants were informed that no additional task would follow and that they could relax.

##### 1.2. Placebo-TSST

The placebo-TSST resembled the TSST but without the committee, the camera, and the microphone: participants were accompanied to the testing room and instructed by the experimenter to sit for 5 minutes and take notes about their career aims, which they would afterwards talk about while standing alone in the room. Following this, they would be asked to perform – for 5 minutes – a simple mental arithmetic task, counting upwards from zero in increments of 15. After the instructions, participants could ask questions about the procedure.

Following the instructions, the experimenter left the room. After 5 minutes, the experimenter entered the room for sampling T6. After the experimenter had left the room, the participants stood up and talked about what they had prepared before. After 5 minutes, the experimenter entered the room, told the participant to start the arithmetic task, and left again. After another 5 minutes, the experimenter re-entered the room for sampling T7. Then, participants were accompanied back to the MRI area.

##### 1.3. Quality assessment of MP2RAGE images

The CAT12 toolbox automatically checks data quality, using a signal-to-noise ratio (SNR) - like parameter, white matter hyperintensities (WMHs), and bias parameters. The images are then rated according to these parameters from A to D. The noise-to-contrast ratio (NCR) is one parameter automatically extracted by CAT12 during preprocessing similar to the SNR, but accounting for WMHs, that is used to rate the quality of the images. We extracted the NCR from the automatically generated quality report of all images. It was then z-scored within sessions. One participant was excluded because of a below-average (zscore < 3) image quality in both scans, two participants were excluded for showing bad image quality in one of the scans and a below-average difference in image quality between the two scan sessions (zscore < 3). One Participant was excluded because of an incidental medical finding. Using a linear mixed model, we found a significant interaction effect of group and time on image quality (see supplements). Posthoc-Tests showed there was no significant group-difference in image quality at any time point, but revealed a significant change in image quality within the control group. This effect was reduced when including mean frame displacement (mFD), a motion parameter obtained from the neighbouring resting state sequences, into the model. The effect of mFD on image quality rated with NCR was close to statistical significant. We therefore included this parameter in our statistical models to minimize the influence of systematic quality changes on volume estimates.

##### 1.4. Pulsed arterial spin labeling

###### 1.4.1. Acquisition

Cerebral blood flow was measured using the pulsed arterial spin labeling (pASL) sequence from Siemens, which employs a proximal inversion with control for off-resonance effects (PICORE) with a thin slice TI1 periodic saturation (Q2TIPS; Luh et al., 1999). Parameters were a 10 cm labeling slab with a 19 mm gap to the imaging slab, TI1 = 700 ms (begin of periodic saturation pulses after inversion), TI1s = 1975 ms (end of periodic saturation pulses after inversion), and TI2 = 2000 ms (begin of image acquisition after the inversion pulse). Interleaved label and control images of 18 slices (ascending order, thickness 4 mm, slice gap 1mm) were acquired using a gradient-echo EPI readout (matrix 64x64, partial Fourier factor 6/8, FOV 192x192 mm2, TR/TE = 3000/15 ms). Each scan consisted of 50 label/control pairs preceded by the acquisition of a single M0 image.

###### 1.4.2. Preprocessing

Original PASL time series were first realigned with FSL *McFlirt*, then normalized to MNI space with a 2-mm isotropic resolution using SPM12, and finally smoothed with a 3D spatial Gaussian filter of 2-mm FWHM. CBF-values were estimated by the perfusion model (see Equation [5] in Wong et al., 1998) from the M0-scaled pairwise differences between the control and label images. The following model parameters were used for 3T: longitudinal relaxation time of arterial blood (T1b) = 1664 ms, longitudinal relaxation time of brain tissue (T1t) = 1330 ms, inversion efficiency = 0.95, exchange time (Tex) = 1600 ms. The TI2 value fed into the model was increased from slice to slice by the inter-slice acquisition time difference of 40 ms.

###### 1.4.3. Quality assessment

In addition to a CBF map, the preprocessing yielded a map of p-values, derived from paired t-tests comparing the control and the label image. All voxels with p > 0.05 were excluded from the analysis, as this may indicate difficulties with signal strength and data interpretation. To identify images with overall bad image quality, the total number of voxels with p > 0.05 was calculated for each participant and images with a number of those voxels exceeding 3 SD above the sample mean were considered outliers, leading to the exclusion of three participants from further pASL-analyses.

CBF values detected with MRI in human GM are distributed around the average of 40-70 ml/100mg/min (Donahue et al., 2006). We therefore excluded values lower than 20ml/100mg/min and values greater than 100 ml/100mg/min from our analysis. As the pASL sequence does not cover the whole brain, a slab is chosen during data acquisition. In this study, the slab was centered around medial temporal regions that included the hippocampus and the amygdala, our anatomical (pre-hoc) ROIs. Consequently, not all VBM clusters were fully covered. To avoid analysing voxels with few available data, an availability mask (Figure S2A) was created, in which each voxel contained the number of images, for which pASL data was available at this voxel. This mask was thresholded at 100 (of 128 possible images, 2 images for each of the 64 participants after QA), binarized, and compared to the VBM result clusters (see Figure S2B). Only clusters, in which at least 70% of voxels overlapped with the availability mask, were included in the post-hoc analysis of CBF.

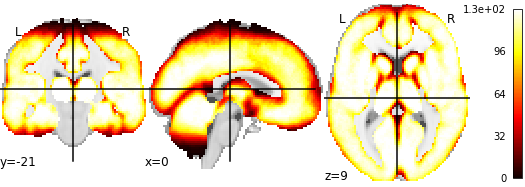

**Figure S2A**: **Availability mask depicting spatial coverage of CBF data from pASL sequence**. The value within each voxel indicates how many images contain usable CBF-data for this voxel (maximum=128). This mask was thresholded at 100 and binarized.

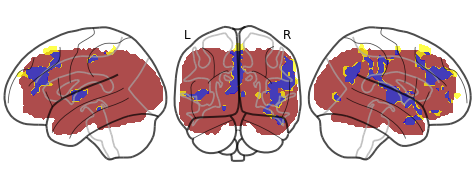

**Figure S2B: VBM-Clusters** (yellow) **included in CBF-analysis (blue)** were chosen based on spatial availability of CBF data (red).

###### 1.4.4. pASL-ROI-Analysis

To test the contribution of blood flow, we investigated CBF changes (measured with pASL) in the main VBM clusters. Five out of 15 VBM clusters were fully covered by the pASL slab (which was chosen to fit the subcortical ROIs, i.e. hippocampus and amygdala). In 8 out of the 15 VBM clusters, including including the three biggest clusters with the strongest statistical effect in the anterior cortical midline and the bilateral insula (the three biggest clusters with the strongest statistical effect), pASL data was available in more than 70% of the voxels. Those clusters were then included in the ROI-based CBF analysis. After quality assessment, 64 participants were included in the analysis.

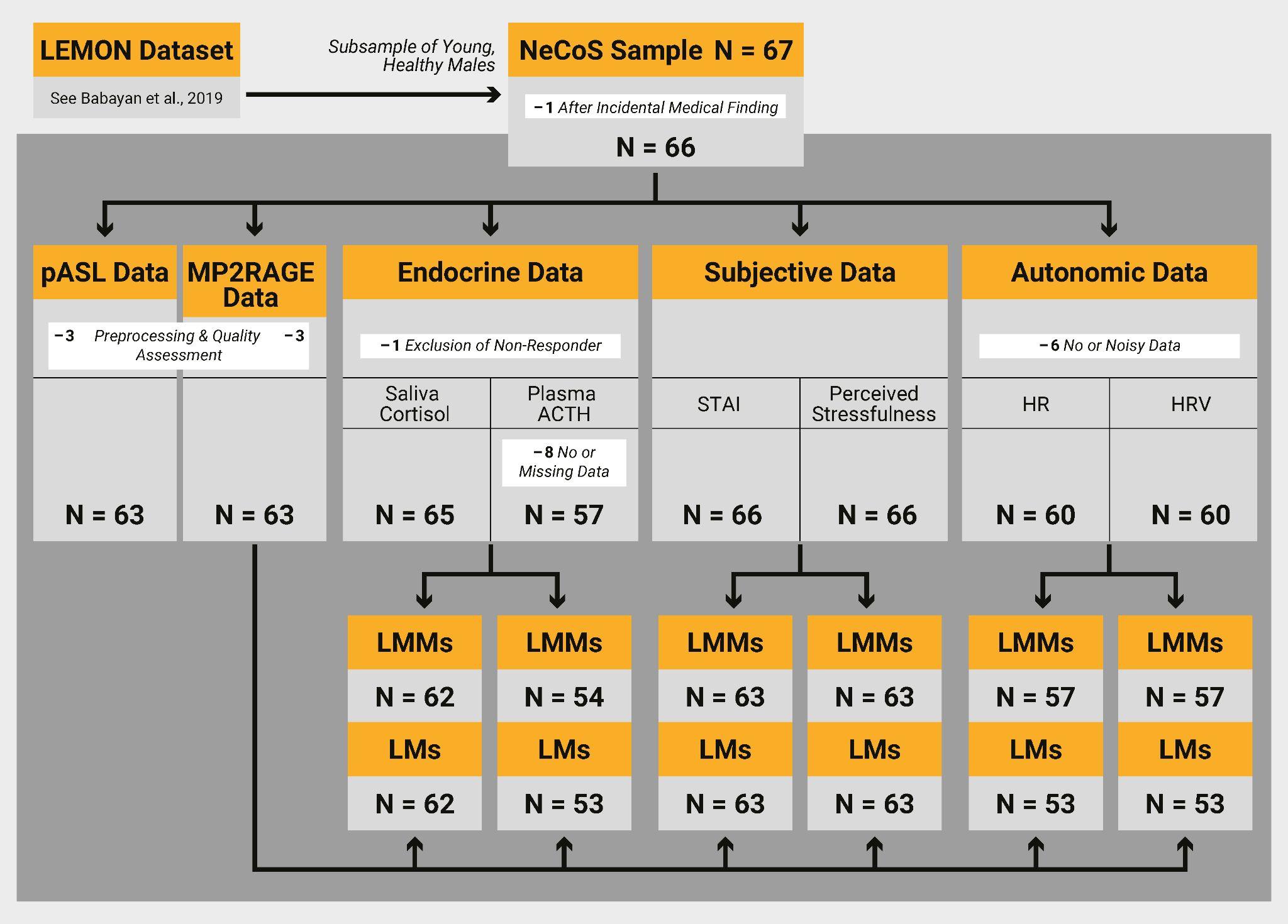

**Figure S3: Flow chart of the acquisition and analysis pipeline with sample sizes at each step.** Abbreviations: LEMON = Sample described in Babayan et al. (2019)

NeCoS = This study. pASL = pulsed arterial spin labeling; L(M)M = linear (mixed) model; HR(V) = heart rate (variability); STAI = state anxiety; ACTH = adrenocorticotropic hormone; Missing data = single samples missing; no data = no data available for this participant.

#### 2. Results

##### 2.1. VBM

###### 2.1.1. Whole-Brain Results

###### 2.1.1.1. 8mm Smoothing Kernel

|  | **Hemis-**  **phere** | **Cluster Name** | **Cluster**  **Size** | **p_FWE_ (TFCE)** | **x y z** | **Linear Mixed Model**  **(Holm-**  **Bonferroni- corrected)** | **Post-Hoc Test (Holm-Bonferroni-corrected)** | **Estimates**  **[Pattern]** |
| --- | --- | --- | --- | --- | --- | --- | --- | --- |
| 1 | L | Superior Medial Frontal Gyrus | 2364 | 0.014 | -03 50 30 | Χ²(1) = 9.1, p = 0.0250 | t/t_c_(66.5) = -4.2,  p = 0.0002  t/t_s_(68.6) = 0.7  p = 0.4915 | $\beta_{c}$ = -0.013*  $\beta_{s}$ = 0.002    [1] |
| 2 | R | (posterior) Insula | 2441 | 0.015 | -12 04 | Χ²(1) = 14.3, p = 0.0019 | t/t_c_(66.5) = - 3.8,  p = 0.0007  t/t_s_(68.8) =   2.2,  p = 0.034 | $\beta_{c}$ = -0.011*  $\beta_{s}$ = 0.007*    [3] |
| 3 | L | (anterior) Insula | 466 | 0.027 | -40 -08 06 | Χ²(1) = 11.3, p = 0.0062 | t/t_c_(66.5) = - 4.3, p = 0.0001  t/t_s_(68.3) = 1.0  p = 0.3012 | $\beta_{c}$ = -0.014*  $\beta_{s}$ = 0.004    [1] |
| 4 | R | Angular Gyrus | 696 | 0.035 | 54 -63 28 | Χ²(1) = 15.5, p = 0.0011 | t/t_c_(66.6) =   0.0,  p = 0.9845  t/t_s_(70.8) = 5.6,  p < 0.0001 | $\beta_{c}$ = 0.0001  $\beta_{s}$ = 0.038*    [2] |
| 5 | L | Parahippocampal Gyrus | 35 | 0.038 | 43 -12 04 | Χ²(1) = 17.4, p = 0.0005 | t/t_c_(66.5) =   4.4,  p < 0.0001  t/t_s_(68.7) =   0.3, p = 0.734 | $\beta_{c}$ = -0.029*  $\beta_{s}$ = -0.002    [1] |
| 6 | R | Inferior Occipital Gyrus | 118 | 0.042 | 52 -80 03 | Χ²(1) = 12.6, p = 0.0042 | t/t_c_(66.6) =   -3.8,  p = 0.0006  t/t_s_(71.2) =   1.7, p = 0.0919 | $\beta_{c}$ = -0.029*  $\beta_{s}$ = 0.015    [1] |
| 7 | L | Mid- Cingulate Cortex | 160 | 0.042 | -03 -24 46 | Χ²(1) = 8.2, p = 0.0214 | t/t_c_(66.5) =   0.4,  p = 0.7114  t/t_s_(68.5) =   4.4,  p = 0.0001 | $\beta_{c}$ = 0.0001  $\beta_{s}$ = 0.014*    [2] |
| 8 | R | Cerebro- Motor- Area | 77 | 0.043 | -04 -06 57 | Χ²(1) = 8.1, p = 0.0214 | t/t_c_(66.8) =   -1.5,  p = 0.1307  t/t_s_(67.5) =   2.7,  p = 0.0193 | $\beta_{c}$ = -0.005  $\beta_{s}$ = 0.009*    [2] |
| 9 | R | Lateral Orbital Gyrus | 79 | 0.045 | 36 39 -15 | Χ²(1) = 15.6, p = 0.0011 | t/t_c_(66.5) =   -3.3,  p = 0.0031  t/t_s_(69.8) =   2.7,  p = 0.0091 | $\beta_{c}$ = -0.020*  $\beta_{s}$ = 0.019*    [3] |
| 10 | R | Precuneus | 141 | 0.046 | 03 -50 57 | Χ²(1) = 11.8, p = 0.0052 | t/t_c_(66.4) =   0.4,  p = 0.7201  t/t_s_(67.7) =   5.2,  p < 0.0001 | $\beta_{c}$ = 0.002  $\beta_{s}$ = 0.026*    [2] |
| 11 | R | Frontal Pole | 52 | 0.046 | 24 63 03 | Χ²(1) = 8.9, p = 0.0175 | t/t_c_(66.4) = - 4.8, p < 0.0001  t/t_s_(68.3) = -0.05,  p = 0.9626 | $\beta_{c}$ = -0.021*  $\beta_{s}$ = -0.0002    [1] |
| 12 | R | Superior Medial Frontal Gyrus | 44 | 0.047 | 06 52 04 | Χ²(1) = 16.6, p = 0.0007 | t/t_c_(67.0) =   -7.5,  p < 0.0001  t/t_s_(67.4) = -0.5,  p = 0.5824 | $\beta_{c}$ = -0.019*  $\beta_{s}$ = -0.002    [1] |
| 13 | L | Superior Temporal Gyrus | 48 | 0.048 | -63 -10 04 | Χ²(1) = 12.4, p = 0.0042 | t/t_c_(66.5) = - 2.9,  p = 0.0088  t/t_s_(68.9) = 2.4,  p = 0.0193 | $\beta_{c}$ = -0.018*  $\beta_{s}$ = 0.017*    [3] |
| 14 |  | // | 1 | 0.048 | 51 -46 51 | – | – | – |
| 15 | R | Middle Frontal Gyrus | 22 | 0.048 | 45 46 12 | Χ²(1) = 8.1, p = 0.0215 | t/t_c_(66.4) =   -3.8,  p = 0.0007  t/t_s_(68.1) =   0.6,  p = 0.5344 | $\beta_{c}$ = -0.016*  $\beta_{s}$ = 0.003    [1] |
| 16 | R | Middle Frontal Gyrus | 23 | 0.050 | 42 54 -02 | Χ²(1) = 7.8, p = 0.0215 | t/t_c_(66.7) =   -3.7,  p < 0.0009  t/t_s_(71.5) =   0.7,  p = 0.5177 | $\beta_{c}$ = -0.027*  $\beta_{s}$ = 0.005    [1] |

**Table S1**. Results from the voxel-based morphometry (VBM) analysis and post-hoc linear mixed models (LMMs) on grey matter volume (GMV) in the VBM clusters. Depicted are hemisphere, cluster name (derived from CAT12’s “neuromorphometrics atlas”), cluster size in voxels, p_FWE_ after threshold-free cluster enhancement (TFCE) correction, coordinates in MNI space (x y z), LMM results, post-hoc-test results, group estimates, and pattern (see Figure 2). The cluster with an extent of 1 voxel was excluded from further analyses. LMM statistical parameters (degrees of freedom (DF), X² value, p value) were obtained from a full-null model comparison. Post-hoc results include the t-ratio with DF and p value. P < 0.05 indicates a significant group-by-time interaction effect. N = 63 (stress group: n = 30).

###### 2.1.1.2. 6mm Smoothing Kernel

To estimate the influence of the smoothing kernel on our results, we repeated the analysis with a 6-mm smoothing kernel. Decreasing the smoothing kernel to 6mm isotropic (FWHM) results in overall fewer and smaller clusters within the same main result regions as with the 8-mm smoothing kernel.

| *Cluster name* | *k* | *p(FWE)* | *Coordinates* |
| --- | --- | --- | --- |
| *Right Posterior Insula* | *462* | *0.012*  *0.035*  *0.0346* | *42 -12 4*  *44 2 2*  *46 6 -9* |
| *Right Superior Medial Frontal Gyrus* | *891* | *0.023*  *0.024*  *0.025* | *2 39 28*  *-3 50 30*  *2 30 27* |
| *Left Anterior Insula* | *191* | *0.027* | *-40 -6 8* |
| *Left Superior Medial Frontal Gyrus* | *164* | *0.034*  *0.046* | *-2 32 56*  *-3 15 50* |
| *Right Central Operculum* | *233* | *0.039*  *0.040*  *0.042* | *56 -20 15*  *39 -24 16*  *60 -9 9* |
| *Left Mid-Cingulate Gyrus* | *75* | *0.041*  *0.041* | *-3 -26 46*  *0 -18 42* |
| *Right Supramarginal Gyrus* | *27* | *0.045* | *63 -24 39* |
| *Left Anterior Cingulate Cortex* | *6* | *0.046* | *-3 42 10* |
| *Left Anterior Insula* | *2* | *0.050* | *-34 2 8* |
| *Right Anterior Insula* | *1* | *0.050* | *40 14 -3* |

**Table S2.** Results from the voxel-based morphometry (VBM) analysis with a 6mm smoothing kernel. Depicted are cluster name (derived from CAT12’s “neuromorphometrics atlas”), cluster size in voxels, p_FWE_ after threshold-free cluster enhancement (TFCE) correction and coordinates in MNI space (x y z).

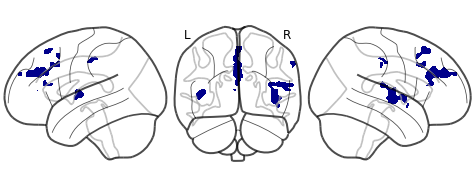

**Figure S4.** Results from Voxel-based morphometry (VBM) on segmented GM images smoothed with 6mm gaussian kernel. Blue clusters indicate a significant (p_FWE_ < 0.05) group-by-time interaction effect on grey matter volume (GMV).

###### 2.1.2. VBM changes in Thalamus cluster from Reinelt & Uhlig et al., (2019)

We analyzed GMV changes in the thalamus cluster from our previous manuscript (Reinelt & Uhlig et al., 2019), which showed a significant group-by-time interaction effect on Eigenvector Centrality values, driven by an increase in EC in the stress group. In this thalamus cluster we find no significant group-by-time interaction effect on GMV (X²(1)=1.66, p=0.1982), but a significant decrease in GMV across both groups (X²(1)=20.42, p<0.0001).

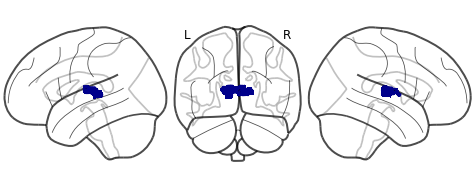

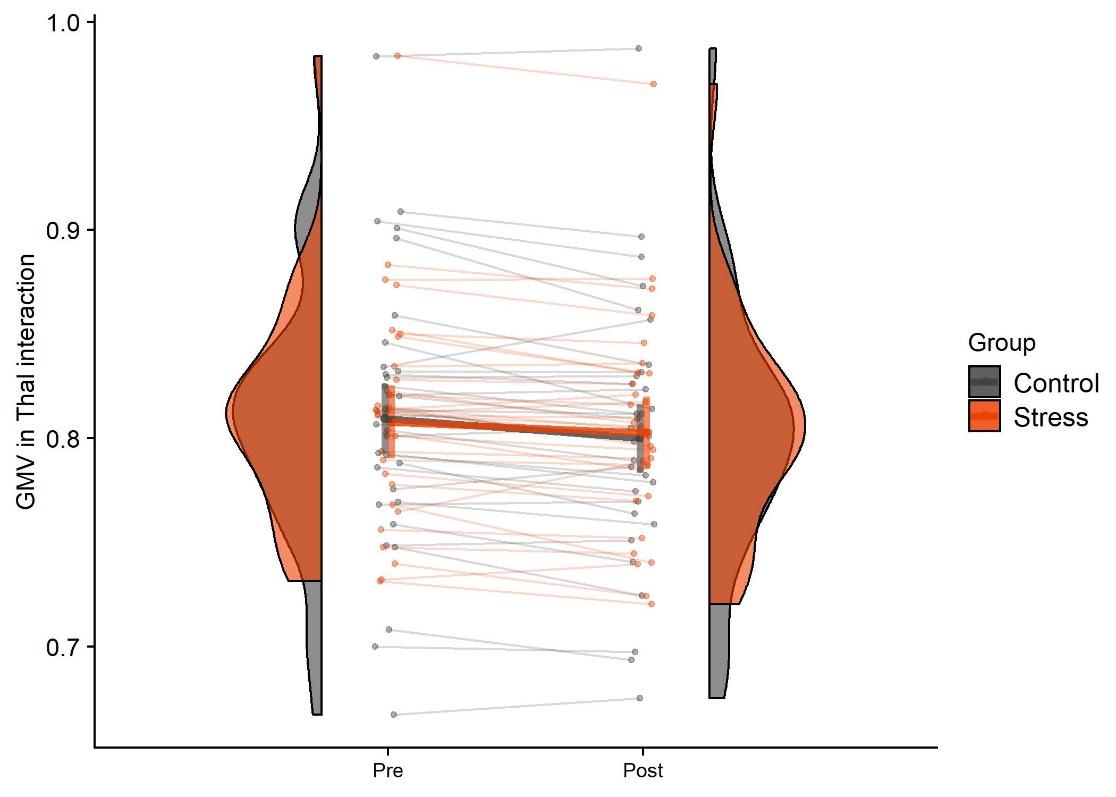

**Figure S5.** Top: ECM clusters (blue) with significant interaction effect of group and time on eigenvector centrality as identified by (Reinelt & Uhlig et al., 2019), Bottom: Extracted GMV values from VBM-analysis in significant interaction cluster from ECM-analysis. There was no significant group-by-time interaction effect for grey matter volume (GMV) but a significant main effect of time (pre > post stress).

###### 2.1.3. Association of VBM changes with other stress measures

| **TEST** | **Variable** | **p(uncor)** | **p(cor)** |
| --- | --- | --- | --- |
| LMM | SalCort | 0.8752 | 1 |
| LMM | ACTH | 0.9797 | 1 |
| LMM | HR | 0.3277 | 1 |
| LMM | HRV | 0.8141 | 1 |
| LMM | STAI | 0.7533 | 1 |
| LMM | VAS | 0.9588 | 1 |
| LM | SalCort | 0.3482 | 1 |
| LM | ACTH | 0.1315 | 1 |
| LM | HR | 0.0003 | 0.0038 |
| LM | HRV | 0.0014 | 0.0149 |
| LM | STAI | 0.0005 | 0.0059 |
| LM | VAS | 0.0210 | 0.1894 |

**Table S3. Association of grey matter volume (GMV) with stress measures.** Results from Linear Mixed Models (LMMs) and Linear Models (LMs) investigating the association of grey matter volume (GMV) and Δ GMV (post-pre) with stress measures and peak reactivity of stress measures. Depicted are Test type, stress measure and p-values as uncorrected and corrected (Bonferroni-Holm) results.

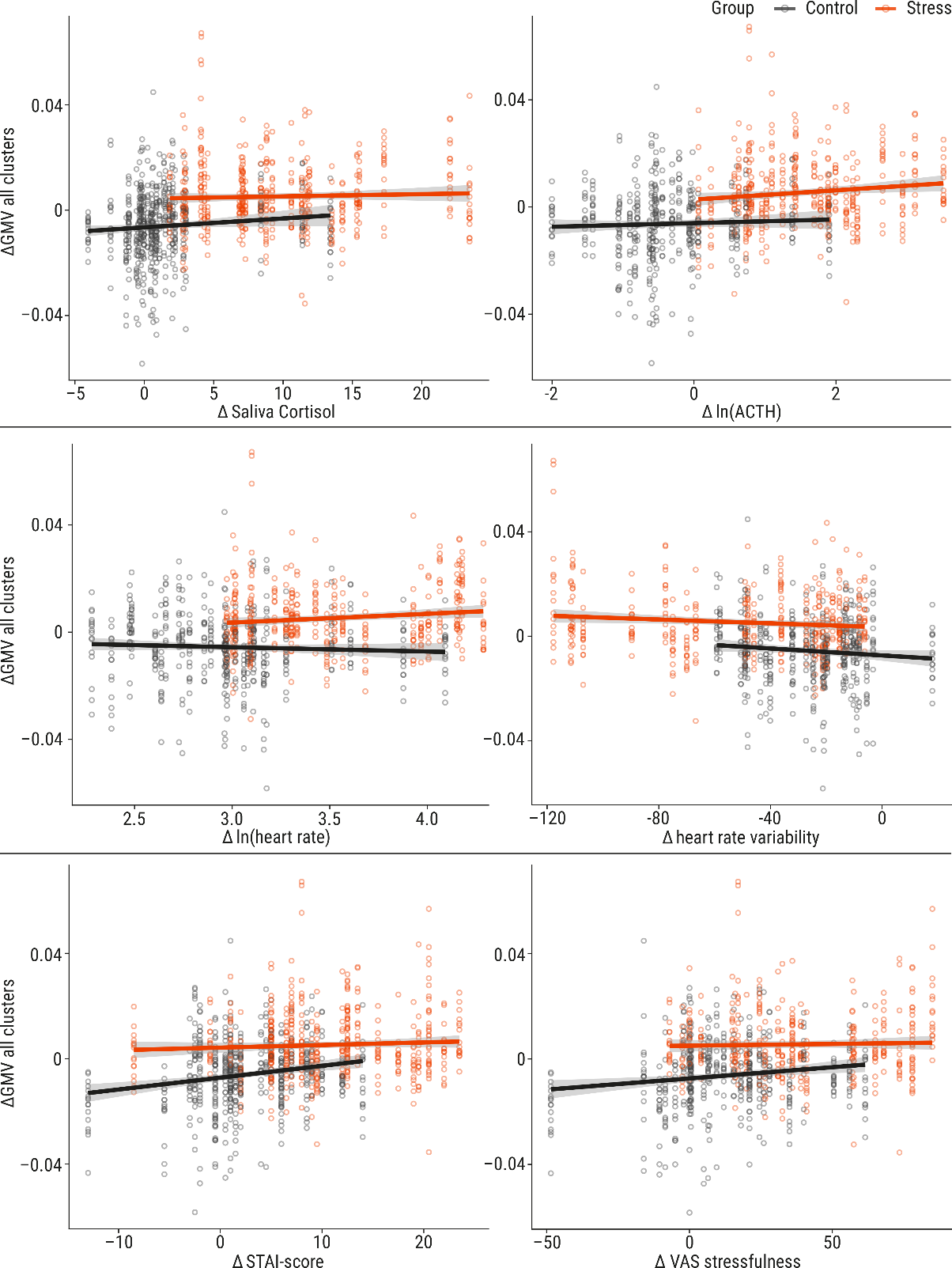

**Figure S6. Association of Δ grey matter volume (GMV; post-pre) with peak reactivity of stress measures.** Shown are significant associations from linear models (LMs): state anxiety (State Anxiety Inventory, STAI; positive association) and heart rate variability (RMSSD; negative association) as well as non-significant association with saliva cortisol, ACTH, heart rate and subjective stressfulness. Line indicates slope and standard error. Points indicate GMV values per voxel-based morphometry (VBM) cluster and subject, each subject is represented in one column of points. Grey: control group, orange: stress group.

### 2.2. T1

| Cluster | emmean T1 in ms (pre,post) | estimate | SE | df | t/t | p-value |
| --- | --- | --- | --- | --- | --- | --- |
| Left Superior Medial Frontal Gyrus | 1651, 1681 | 35.18 | 16.9 | 1812 | 2.080 | 0.3010 |
| Right Posterior Insula | 1501, 1512 | 11.17 | 16.9 | 1812 | 0.661 | 1.0000 |
| Left Anterior Insula | 1637, 1640 | 3.03 | 16.9 | 1812 | 0.179 | 1.0000 |
| Right Angular Gyrus | 1003, 1087 | 83.95 | 16.9 | 1812 | 4.965 | <0.0001 |
| Left Parahippocampal Gyrus | 1437,  1441 | 3.32 | 16.9 | 1812 | 0.196 | 1.0000 |
| Right Inferior Occipital Gyrus | 705, 799 | 94.37 | 17.1 | 1812 | 5.508 | <0.0001 |
| Left Mid-Cingulate Cortex | 1617,  1612 | -5.18 | 16.9 | 1812 | -0.306 | 1.0000 |
| Right Cerebromotor Area | 1558, 1554 | -4.08 | 16.9 | 1812 | -0.241 | 1.0000 |
| Right Lateral Orbital Gyrus | 1205, 1335 | 130.73 | 16.9 | 1812 | 7.731 | <0.0001 |
| Left Precuneus | 1760, 1756 | 4.48 | 16.9 | 1812 | -0.265 | 1.0000 |
| Right Frontal Pole | 1384,  1481 | 97.03 | 16.9 | 1812 | 5.738 | <0.0001 |
| Right Superior Medial Frontal Gyrus | 1639, 1646 | 7.16 | 16.9 | 1812 | 0.423 | 1.0000 |
| Left Superior Temporal Gyrus | 1239, 1301 | 61.94 | 16.9 | 1812 | 3.663 | 0.0026 |
| Right Middle Frontal Gyrus | 1376, 1431 | 55.37 | 16.9 | 1812 | 3.275 | 0.0097 |
| Right Middle Frontal Gyrus 2 | 977, 1127 | 150.01 | 16.9 | 1812 | 8.871 | <0.0001 |

**Table S4. Post-Hoc Tests of T1 values in VBM clusters**. Depicted are T1 values per cluster and time point based on estimated marginal means (emmeans), degrees of freedom (df), standard error (SE), t-ratio and p-value.

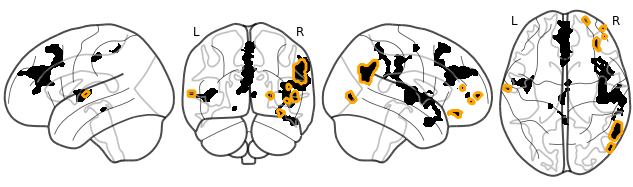

**Figure S7.** VBM clusters (black) with significant increase in T1 values (orange) across groups.

**Increase in overall GM at different thresholds of GM probability**

| **Timepoint** | **GM Threshold** | **emmean** | **lower.CL** | **upper.CL** | **Increase in ms** | **Post-Hoc Test (Bonferroni-Holm)** |
| --- | --- | --- | --- | --- | --- | --- |
| Pre | 01 | 1209 | 1195 | 1222 | 28.79 | t/t=8.974, p<.0001 |
| Post | 01 | 1237 | 1224 | 1251 |  |  |
| Pre | 02 | 1311 | 1298 | 1325 | 24.44 | t/t=7.619, p<0.0001 |
| Post | 02 | 1336 | 1322 | 1349 |  |  |
| Pre | 03 | 1369 | 1356 | 1383 | 19.40 | t/t=6.047, p<0.0001 |
| Post | 03 | 1388 | 1375 | 1402 |  |  |
| Pre | 05 | 1409 | 1396 | 1423 | 1.85 | t/t=0.577, p =0.5645 |
| Post | 05 | 1411 | 1397 | 1424 |  |  |

**Table S5 Results from linear mixed models (LMMs) on T1 values in GM at different tissue probability thresholds.** Depicted are Timepoint, Threshold, estimated marginal mean, upper and lower confidence interval (95%), increase in milliseconds (ms), results from post-hoc tests (within group, t-ratio and p-value adjusted for multiple comparisons with Bonferroni-Holm method).

##### 2.3. T1w intensity

While we found neither significant group-by-time interaction effect on T1w intensity values (Χ²(1) = 3.13, p = 0.99, Figure 4), nor a significant change over time across groups (X²(1) = 2.16, p = 0.141), we identified a significant main effect of group (X²(1) = 6.9, p = 0.0087). To further follow up on this, we investigated baseline differences and potentially contributing parameters: T1w intensity values showed significant group-differences at baseline (t/tpre(78.4) = -2.35, p = 0.0178), which persisted until the post-intervention scan (t/tpre(77.3) = -2.53, p = 0.0212). These group-differences in signal intensity may arise from inter-individual differences, such as TIV, head motion, or other characteristics that we did not record (e.g., body temperature):

1.	For TIV, there was no significant main effect of group (X²(1) = 0.11, p = 0.7452).

2.	Head motion was approximated by the mean framewise displacement (MFD) in the resting-state scan directly preceding the MP2RAGE scans. There was no significant main effect of group on MFD (X²(1) = 0.08, p = 0.7793).

In LMMs, neither TIV (X²(1) = 0.77, p = 0.3803) nor MFD (X²(1) = 0.94, p = 0.3335) showed a significant effect on T1w intensity values. In an additional control analysis, we found that the significant group difference in T1w intensity values was also present in WM (F(1) = 5.16, p = 0.02667). No other variables (CBF: t/tpre(964) = 1.57, p = 0.2341, GMV: t/tpre(85.5) = 1.26, p = 0.4216, T1: t/tpre(82.4) = 1.16, p = 0.2511) showed a significant main effect of group in the pre-intervention scan (i.e., baseline differences).

##### 2.4. PASL

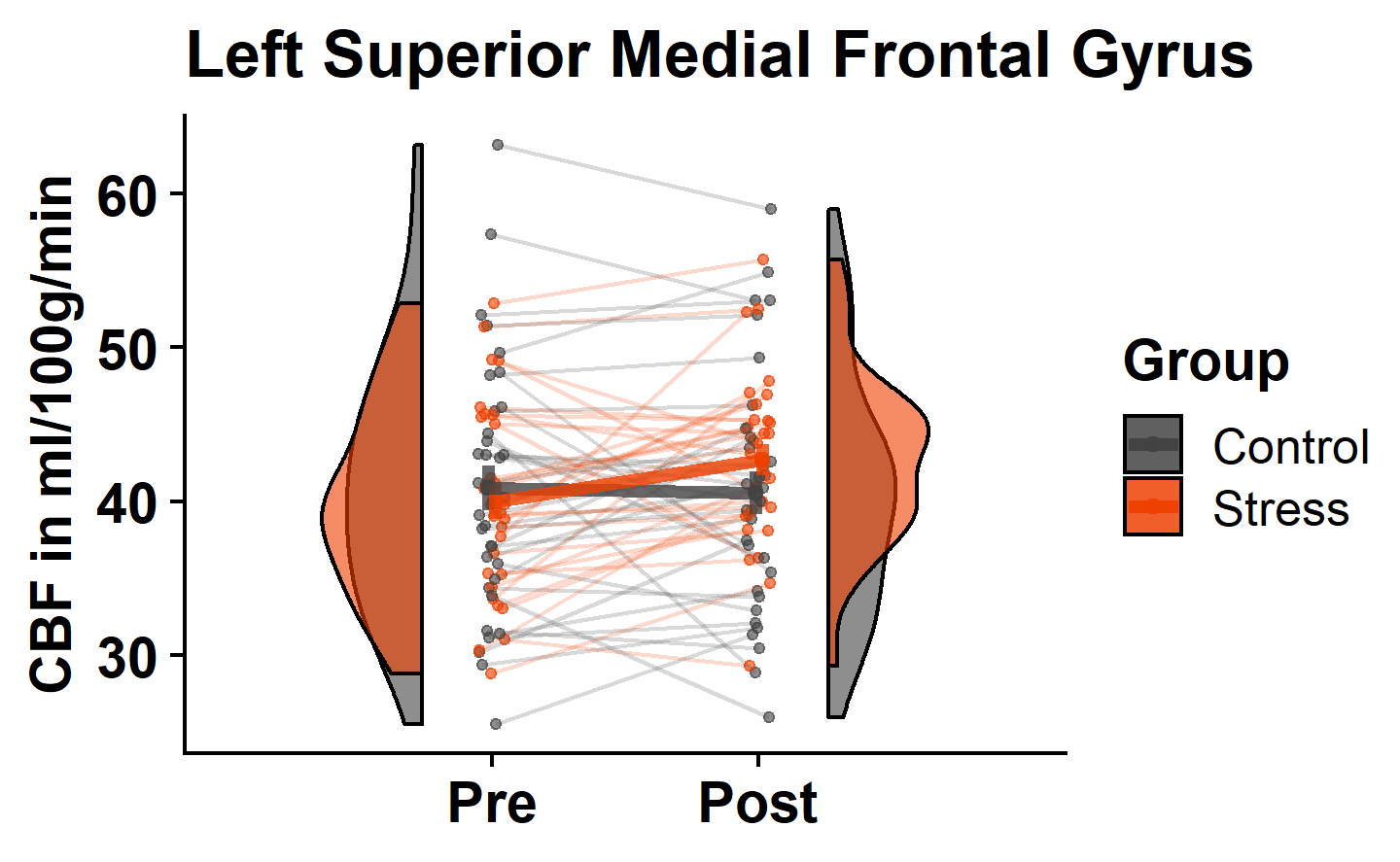

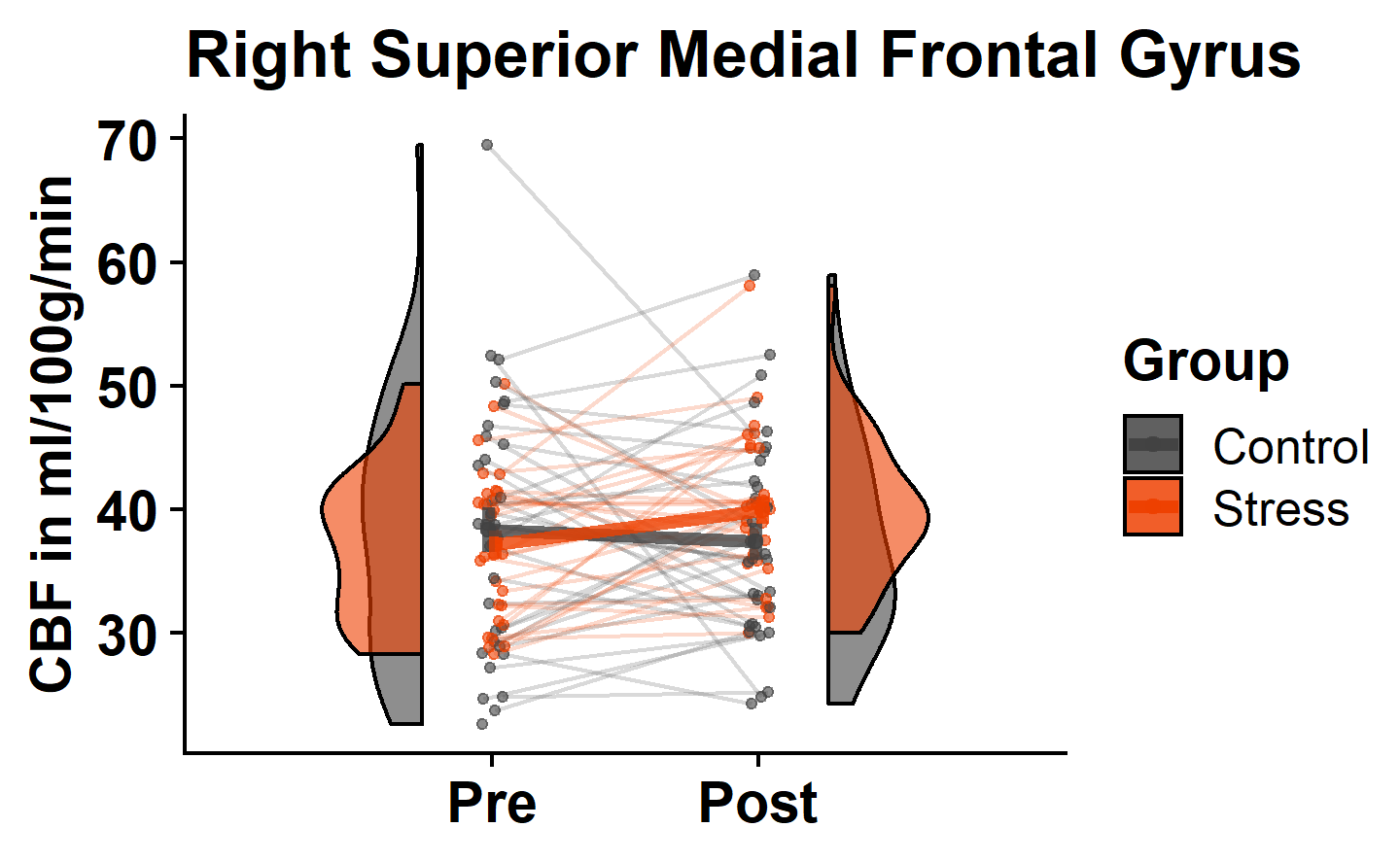

**Figure S8**. **CBF changes in the Left and Right Superior Medial Frontal Gyrus (SMFG)** in the control group (grey) and in the stress group (red). Shown are scans (points) per subject (thin lines) averaged across clusters and group distributions (half violin) for pre- and post-scan. Bold lines indicate estimated marginal means and 95% confidence intervals obtained from linear mixed models. If data were transformed (ln or square-root) for statistical analysis, the estimates were back-transformed for visualization.

In a control analysis, we found that neither age (X²(1) = 0.59, p = 0.4455) nor TIV (X²(1) = 0.01, p = 0.916) nor MFD (X²(1) = 0.01, p = 0.9060) significantly influences CBF values so we did not include them as covariates.

| **Hemisphere** | **Clustername** | **xyz** | **pFWE(TFCE)** | **Test** | **N** | **DF** | **Stat** | **p_uncor** | **p_cor** | **Estimate** |
| --- | --- | --- | --- | --- | --- | --- | --- | --- | --- | --- |
| L | Superior Medial Frontal Gyrus | -03 50 30 | 0.014 | Interaction | 63 | 1 | 4.51 | 0.0338 | 0.2701 | NA |
|  |  |  |  | Main Effect Time | 63 | 1 | 2.68 | 0.1018 | 0.8142 | NA |
|  |  |  |  | Posthoc Control | 33 | 65 | -0.25 | NA | 0.8057 | -0.0061 |
|  |  |  |  | Posthoc Stress | 30 | 65 | 2.7 | NA | 0.0175 | 0.0702 |
| R | Posterior Insula | 43 -12 04 | 0.015 | Interaction | 63 | 1 | 0.05 | 0.8271 | 1 | NA |
|  |  |  |  | Main Effect Time | 63 | 1 | 1.5 | 0.2204 | 1 | NA |
|  |  |  |  | Posthoc Control | 33 | 65 | 1.03 | NA | 0.6168 | 0.023 |
|  |  |  |  | Posthoc Stress | 30 | 65 | 0.68 | NA | 0.6168 | 0.016 |
| L | Anterior Insula | -40 -08 06 | 0.027 | Interaction | 63 | 1 | 0.4 | 0.5287 | 1 | NA |
|  |  |  |  | Main Effect Time | 63 | 1 | 0.44 | 0.5088 | 1 | NA |
|  |  |  |  | Posthoc Control | 33 | 65 | 0.9 | NA | 0.7415 | 0.0277 |
|  |  |  |  | Posthoc Stress | 30 | 65 | 0 | NA | 0.9988 | 0 |
| L | Parahippocampal Gyrus | 43 -12 04 | 0.038 | Interaction | 63 | 1 | 0 | 0.9954 | 1 | NA |
|  |  |  |  | Main Effect Time | 63 | 1 | 2.33 | 0.1266 | 0.886 | NA |
|  |  |  |  | Posthoc Control | 32 | 64 | 1.11 | NA | 0.5417 | 0.0447 |
|  |  |  |  | Posthoc Stress | 30 | 64 | 1.05 | NA | 0.5417 | 0.0443 |
| L | Midcingulate Cortex | -03 -24 46 | 0.042 | Interaction | 63 | 1 | 0.61 | 0.4348 | 1 | NA |
|  |  |  |  | Main Effect Time | 63 | 1 | 0.5 | 0.4778 | 1 | NA |
|  |  |  |  | Posthoc Control | 32 | 65 | 0.03 | NA | 0.9723 | 0.0012 |
|  |  |  |  | Posthoc Stress | 30 | 63 | -1.04 | NA | 0.6024 | -0.0368 |
| R | Superior Medial Frontal Gyrus | 06 52 04 | 0.047 | Interaction | 63 | 1 | 3.17 | 0.0748 | 0.5235 | NA |
|  |  |  |  | Main Effect Time | 63 | 1 | 0.87 | 0.3498 | 1 | NA |
|  |  |  |  | Posthoc Control | 32 | 64 | 0.55 | NA | 0.5863 | 0.0206 |
|  |  |  |  | Posthoc Stress | 30 | 64 | -1.94 | NA | 0.113 | -0.0769 |
| R | Middle Frontal Gyrus | 45 46 12 | 0.048 | Interaction | 63 | 1 | 0.26 | 0.6105 | 1 | NA |
|  |  |  |  | Main Effect Time | 63 | 1 | 0 | 0.9876 | 1 | NA |
|  |  |  |  | Posthoc Control | 32 | 65 | 0.34 | NA | 1 | 0.0133 |
|  |  |  |  | Posthoc Stress | 30 | 64 | -0.37 | NA | 1 | -0.0151 |
| R | Middle Frontal Gyrus | 42 54 -02 | 0.05 | Interaction | 63 | 1 | 0.26 | 0.6105 | 1 | NA |
|  |  |  |  | Main Effect Time | 63 | 1 | 0 | 0.9876 | 1 | NA |
|  |  |  |  | Posthoc Control | 32 | 65 | 0.34 | NA | 1 | 0.0133 |
|  |  |  |  | Posthoc Stress | 30 | 64 | -0.37 | NA | 1 | -0.0151 |

**Table S6. Results from linear mixed models (LMMs) on CBF values in VBM clusters.** Depicted are hemisphere, cluster name (derived from CAT12’s “neuromorphometrics atlas”), results from LMM (group-by-time interaction effect & main effect of time) obtained from a full-null model comparison, and post-hoc test (estimated marginal means). Statistical parameters include degrees of freedom (DF), sample size (N), Test Statistic (Stat: X² for LMM and t-ratio for posthoc test), as well as p-value (uncorrected and corrected with Holm method) and estimate). P<0.05 indicates statistical significance.

| **Clustername** | **scan** | **stress** | **emmean** | **SE** | **lower CL** | **upper CL** |
| --- | --- | --- | --- | --- | --- | --- |
| Left Superior Medial Frontal Gyrus | pre | 0 | 40.87 | 1.49 | 37.93 | 43.8 |
|  | post | 0 | 40.56 | 1.49 | 37.63 | 43.5 |
|  | pre | 1 | 39.98 | 1.56 | 36.9 | 43.06 |
|  | post | 1 | 42.73 | 1.56 | 39.65 | 45.81 |
| Left Anterior Insula | pre | 0 | 39.55 | 1.49 | 36.62 | 42.49 |
|  | post | 0 | 38.39 | 1.49 | 35.46 | 41.33 |
|  | pre | 1 | 40.67 | 1.56 | 37.59 | 43.75 |
|  | post | 1 | 40.29 | 1.56 | 37.21 | 43.37 |
| Right Posterior Insula | pre | 0 | 41.01 | 1.49 | 38.07 | 43.94 |
|  | post | 0 | 40.06 | 1.49 | 37.13 | 43 |
|  | pre | 1 | 41.68 | 1.56 | 38.6 | 44.76 |
|  | post | 1 | 40.91 | 1.56 | 37.83 | 43.99 |
| Left Parahippocampal Gyrus | pre | 0 | 44.69 | 1.5 | 41.72 | 47.65 |
|  | post | 0 | 43.08 | 1.49 | 40.15 | 46.02 |
|  | pre | 1 | 44.9 | 1.58 | 41.79 | 48.01 |
|  | post | 1 | 43.17 | 1.56 | 40.09 | 46.25 |
| Left Midcingulate Cortex | pre | 0 | 42.37 | 1.5 | 39.4 | 45.33 |
|  | post | 0 | 42.44 | 1.5 | 39.47 | 45.4 |
|  | pre | 1 | 42.35 | 1.56 | 39.27 | 45.43 |
|  | post | 1 | 43.88 | 1.56 | 40.8 | 46.95 |
| Right Superior Medial Frontal Gyrus | pre | 0 | 38.33 | 1.49 | 35.39 | 41.27 |
|  | post | 0 | 37.38 | 1.5 | 34.42 | 40.34 |
|  | pre | 1 | 36.82 | 1.58 | 33.71 | 39.93 |
|  | post | 1 | 39.94 | 1.56 | 36.86 | 43.02 |
| Right Middle Frontal Gyrus | pre | 0 | 40.71 | 1.5 | 37.74 | 43.67 |
|  | post | 0 | 40.04 | 1.49 | 37.11 | 42.98 |
|  | pre | 1 | 40.53 | 1.56 | 37.45 | 43.6 |
|  | post | 1 | 40.52 | 1.56 | 37.44 | 43.6 |
| Right Middle Frontal Gyrus 2 | pre | 0 | 45.39 | 1.49 | 42.45 | 48.32 |
|  | post | 0 | 43.91 | 1.52 | 40.92 | 46.9 |
|  | pre | 1 | 45.28 | 1.61 | 42.1 | 48.46 |
|  | post | 1 | 46.83 | 1.56 | 43.75 | 49.91 |

**Table S7. Estimated Marginal means of CBFvalues in VBM clusters**. Depicted are CBF-values per cluster, time point and group based on estimated marginal means (emmeans), standard error (SE), upper and lower 95%-confidence level (CL).

##### 2.5. Overview Tables

| *Statistical Test* | | *VBM* | *Total Tissue Volumes (VBM)*  *GM WM CSF* | | | *CBF* | *T1 values* | *T1w*  *intensity values* |
| --- | --- | --- | --- | --- | --- | --- | --- | --- |
| *Global Interaction Effect* | | **interaction*** | **interaction*** | n.s. | **interaction*** | **interaction*** | n.s. | n.s. |
| *Direction of change*  *(* indicates p<0.05)* | *Stress Group* | increase | increase | —------- | decrease | increase | ---------- | ---------- |
|  | *Control Group* | decrease | decrease | —------- | **increase*** | decrease | ---------- | ---------- |
| *main effect of time* | | **---------------** | **---------------** | n.s. | **---------------** | ---------- | **increase*** | n.s. |

**Table S8. Overview of different modalities (global):** Results from global linear mixed models (LMMs) on GMV, total tissue volumes (from VBM),T1 values, T1-weighted intensity values and CBF values in VBM clusters. Depicted are Modality and Test type cell content reflects the result in words (*indicates p<0.05, “n.s.” indicates no significant effect, — indicates no test was conducted). The direction of change for post-hoc tests after significant LMMs is reported irrespective of p-value.

|  | **Hemis-**  **phere** | **Clustername** | **Cluster-**  **size** | **VBM Effect** | **Pasl Effect** | **T1 Effect** | **UNI Effect** |
| --- | --- | --- | --- | --- | --- | --- | --- |
| 1 | L | Superior Medial Frontal Gyrus | 2364 | **Interaction**  **Control: Decrease**  **Stress: None** | **Interaction**  **Control: None**  **Stress: Increase** | None | None |
| 2 | R | (posterior) Insula | 2441 | **Interaction**  **Control: Decrease**  **Stress: Increase** | None | None | None |
| 3 | L | (anterior) Insula | 466 | **Interaction**  **Control: Decrease**  **Stress: None** | None | None | None |
| 4 | R | Angular Gyrus | 696 | **Interaction**  **Control: None**  **Stress: Increase** | NA | **Main Effect of Time (Increase)** | None |
| 5 | L | Parahippocampal Gyrus | 35 | **Interaction**  **Control: Decrease**  **Stress: None** | **No Interaction**  **Main Effect of Time (Decrease)** | None | None |
| 6 | R | Inferior Occipital Gyrus | 118 | **Interaction**  **Control: Decrease**  **Stress: None** | NA | **Main Effect of Time (Increase)** | None |
| 7 | L | Mid-Cingulate Cortex | 160 | **Interaction**  **Control: None**  **Stress: Increase** | None | None | None |
| 8 | R | Cerebro-Motor- Area | 77 | **Interaction**  **Control: None**  **Stress: Increase** | NA | None | None |
| 9 | R | Lateral Orbital Gyrus | 79 | **Interaction**  **Control: Decrease**  **Stress: Increase** | NA | **Main Effect of Time (Increase)** | None |
| 10 | R | Precuneus | 141 | **Interaction**  **Control: None**  **Stress: Increase** | NA | None | None |
| 11 | R | Frontal Pole | 52 | **Interaction**  **Control: Decrease**  **Stress: None** | NA | **Main Effect of Time (Increase)** | None |
| 12 | R | Superior Medial Frontal Gyrus | 44 | **Interaction**  **Control: Decrease**  **Stress: None** | **Interaction**  **Control: None**  **Stress: Increase** | None | None |
| 13 | L | Superior Temporal Gyrus | 48 | **Interaction**  **Control: Decrease**  **Stress: Increase** | NA | **Main Effect of Time (Increase)** | None |
| 14 |  | NA | 1 | NA | NA |  |  |
| 15 | R | Middle Frontal Gyrus | 22 | **Interaction**  **Control: Decrease**  **Stress: None** | None | **Main Effect of Time (Increase)** | None |
| 16 | R | Middle Frontal Gyrus | 23 | **Interaction**  **Control: Decrease**  **Stress: None** | None | **Main Effect of Time (Increase)** | None |

**Table S9. Overview of different modalities (local):** Results from local linear mixed models (LMMs) on GMV, T1 values, T1-weighted intensity values and CBF values in VBM clusters. Depicted are hemisphere, cluster name (derived from CAT12’s “neuromorphometrics atlas”), cluster size and the significant result (or: “none”) in words (Interaction or main effect and the direction of change). NA indicates no test was conducted.
